# A Neurotensin Brake on Exploratory Drive under Persistent Threat

**DOI:** 10.64898/2026.07.29.741073

**Authors:** Molly McDougle, Victoria Glass, Subramanian Suganya, Whitnei Smith, Paula Frost, Bella DePasquale, Lanah Ha, Alyssa Koehler, Kamryn Thomas, Jack Capurso, Jose H. Ledo, Stefano Berto, Estefania P. Azevedo

## Abstract

Avoidance behavior is an adaptive response that delays exploration to promote survival. Avoidance is enhanced by psychological stress and is a hallmark of many neuropsychiatric disorders. Neural circuits that control avoidance by integrating stressful stimuli and modulating exploratory behavior remain underexplored. Elucidating the functional dynamics of this highly conserved phenomenon and the underlying neural mechanism of avoidance is an important open-ended question, with relevance to understanding both innate behaviors and neuropsychiatric disorders. Using predator odor as an innate, chronic stressor to increase avoidance behaviors in mice, we identified a neural population in the lateral septum (LS) that integrates threat information and modulates latency to explore. Calcium recordings in freely exploring mice combined with activity-based transcriptomics revealed that predator-responsive LS neurons are GABAergic and express neurotensin (LS^NT^). Further, single nuclei RNA-seq analysis revealed that among predator-responsive neurons, NT-enriched inhibitory clusters are predominant. Chronic activation of LS^NT^ neurons induces avoidance behaviors in the absence of predator odor, while synaptic silencing of this population abrogates predator-enhanced avoidance. Using transgenic mouse models to indelibly tag predator-responsive neurons, we defined the downstream circuit that connects the encoding of predator odor information to the lateral hypothalamus. Projection-specific activation of LS^NT^→LHA neurons recapitulate stress-induced avoidance behaviors in mice. Finally, we showed that deletion of neurotensin from LS neurons prevented the effects of predator odor on exploration. Together, these findings offer a genetically-and projection-defined, top-down circuit linking the limbic neurotensinergic system to chronic psychological stress and avoidance behaviors in mice.

## INTRODUCTION

Novelty is an essential aspect of life that drives exploratory behavior in animals(1,2). Exploration in response to novelty is a process by which animals investigate new environments to gather information about resources such as food, water, mating partners, or safe shelter(1,2). This behavior is highly conserved across species and fundamental to survival. However, novelty also presents the potential for danger as unknown foods may be toxic, and unknown environments may harbor predators (1,2). Consequently, various species show fear and subsequent avoidance of novel stimuli(3). Avoidance is an adaptive and protective behavior, increasing hesitance and vigilance before and during exploration of the unknown(3,4). This behavior helps to limit the danger associated with novelty. An appropriate level of avoidance should minimize risk in a context-specific manner. In humans, pathological adaptation to novelty and avoidant behavior has been implicated in many neuropsychiatric disorders, including autism, anxiety disorders, eating disorders, posttraumatic stress disorder (PTSD) and schizophrenia (3,4). Critically, chronic psychological stress enhances avoidance in both humans and rodents (5). In the presence of stressful stimuli, avoidance behaviors (i.e. latency to approach, anxiety-like behaviors, approach/avoidance conflict) are enhanced and can decrease exploration and thus appetitive behaviors in rodents and humans (5–7).

In rodents, chronic psychological stress models include maternal separation, loud noises, immobilization, and predation cues (8). Exposure to predator odor (either natural or synthetic) causes rapid activation of the sympathetic nervous system and HPA axis, leading to the development of anxiety-like behavior(9). As a result, predator-based models have been widely used to model anxiety-like behaviors and avoidance in rodents (9–13). Most studies using predator odors to induce anxiety-like behaviors and avoidance employ synthetic odors derived from the red fox, a natural mouse predator, or its analogs: 2,5-dihydro-2,4,5-trimethylthiazoline (TMT), 2,4,5-trimethylthiazole (12–16) and 2-methyl-2-thiazoline (17,18). These synthetic predator odors are detected through a variety of olfactory receptors (Olfr1019, Olfr30, Olfr376, Olfr20, Olfr1047 and Olfr1079) in the olfactory bulb (OB) that sends olfactory information to the piriform cortex (PC) (19,20). PC neurons project to many limbic regions directly and indirectly, thus rapidly relaying olfactory predator information to brain areas involved in emotional, cognitive, and motor control(21). Limbic structures implicated in the response to physical and psychological stressors include the central amygdala (CeA), bed nucleus of the stria terminalis (BNST), hippocampus and lateral septum (LS) (22). Previous work has shown that neurons in the LS rapidly respond to threats and aversive stimuli to control innate behaviors such as feeding, social behaviors, and escape (23–26). Indeed, the LS is a key node in the limbic system that relays emotional information from the cortex to the hypothalamus and midbrain to control motivated behaviors (22,27). Neurons of the LS are mainly GABAergic, with many subtypes that communicate via local circuits to control many aspects of goal-directed behaviors (23–27). The LS receives sensory information from various cortical areas, such as the medial prefrontal cortex, entorhinal cortex and the piriform cortex, suggesting that the LS may also be relevant to processing many modalities of sensory information, including aversive, threat-signaling odors (23–27). However, the precise neural circuits controlling avoidance behaviors induced by the exposure to predator odors are unknown. Understanding the mechanistic basis of stress-enhanced avoidance can help us better understand the pathophysiology of many neuropsychiatric disorders and fundamental animal behavior.

Here we show that chronic exposure to predator odor (PO) increases avoidance behaviors in mice through specific limbic mechanisms. Using whole brain Fos mapping and unbiased activity-based transcriptomics, we identified a cell type-specific population in the lateral septum (LS) that integrates predator odor information and modulates the latency to explore novel environments, even when animals are fasted and familiar foods are present. We further explored the molecular properties of this population using bioinformatics and single-cell RNAseq analysis, revealing that only a handful of clusters contain the transcripts enriched in response to predator odor. Among these, peptidergic and GABAergic clusters are enriched, including LS neurons that express neurotensin (LS^NT^). Calcium recordings in freely exploring mice revealed that LS^NT^ neurons are responsive to predator odor. Chronic activation of LS^NT^ neurons robustly induced avoidance behaviors in the absence of predator odor, while synaptic silencing of this ensemble completely abrogated predator-enhanced avoidance. Using genetic mouse models to indelibly tag predator odor-responsive neurons, we defined the downstream circuit that connects the encoding of predator odor information to the lateral hypothalamus (LHA). We further show that projection-specific activation of LS^NT^◊LHA neurons control stress-induced avoidance behaviors in mice. Using genetic tools to conditionally delete neurotensin from LS neurons, we show that septal neurotensin release is required for enhanced avoidance behaviors induced by predator odor. Together, these findings offer a genetically-and projection-defined, top-down circuit linking psychological stress to avoidance behaviors in mice via neurotensinergic signaling.

## RESULTS

### Chronic predator odor exposure shifts coping behaviors in mice from exploration to avoidance

In nature, rodents are mostly nocturnal and depend heavily on olfactory cues to detect predators, because vision is limited under low-light conditions and odors linger even when the predator is out of sight (28). Predator odor is a potent natural stressor that elicits avoidance behaviors and activates neuroendocrine pathways critical for survival (17–20). Acute exposure to predator odor triggers the hypothalamic-pituitary-adrenal (HPA) axis, leading to elevated glucocorticoid levels that modulate stress responses (17–20). Repeated exposure to such stressors can lead to attenuation of the HPA axis response, a result of habituation and allostasis (29).

We designed an experiment to measure avoidance behavior following acute or chronic predator odor exposure. To assess the impact of predator odor exposure on avoidance, wild-type mice were exposed to predator odor (PO) in an identical clean cage for 1h (acute exposure) or 1h/day for 30 days (chronic exposure) (Fig.1A). Twenty-four hours after the last exposure, without the presence of PO, the same mice underwent the novelty-suppressed feeding test (NSF, Fig. 1B) and the elevated plus maze test (EPM, Fig. 1C). The NSF test measures the latency to approach a familiar food in a new environment when animals are hungry (overnight fasting) while EPM tests exploration of a new environment (open arms vs closed arms) in the absence of food. Mice exposed for 1h did not exhibit significant changes in avoidance behavior in either task compared to unexposed controls (NSF: p=0.69; EPM (open arms): p=0.26; EPM (closed arms): p=0.058). After 30 days of daily PO exposure, a marked shift emerged, with mice displaying significant avoidance behavior in the NSF (***p=0.0002) and in the EPM test (EPM (open arms): *p=0.02, EPM (closed arms): **p=0.0073). We also observed increased locomotion (distance walked) after 30 days of exposure to PO (*p=0.04), but not after a single exposure (p=0.99). By 30 days, mice exhibited robust avoidance responses across both behavioral paradigms, suggesting that prolonged exposure to predator odor altered exploratory behavior over time, shifting coping strategies from adaptive exploration to maladaptive avoidance.

**Figure 1.**
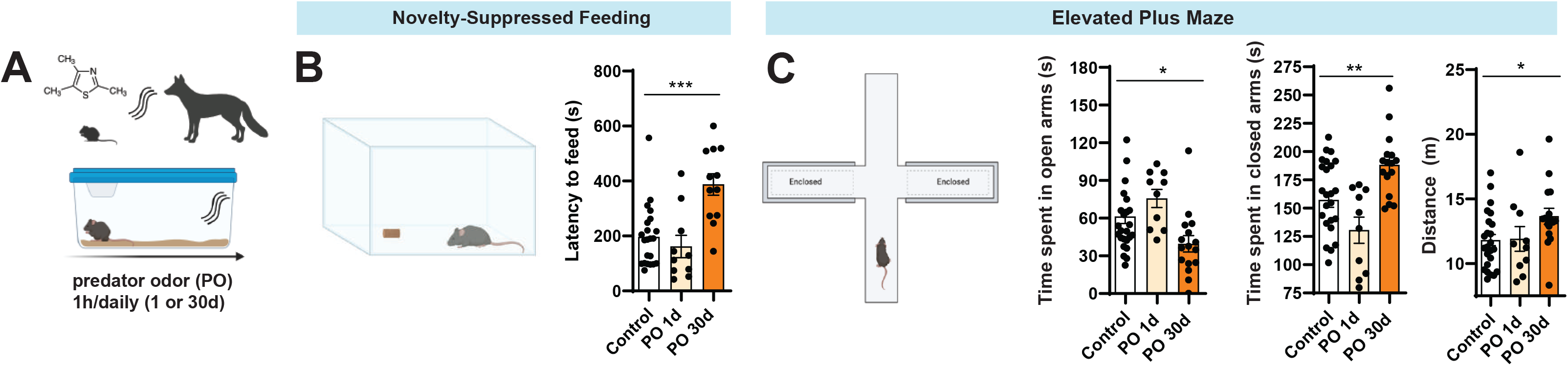
Chronic predator odor exposure increases avoidance behavior in mice. (A) Schematic of the experimental design. Mice were exposed to predator odor (PO) for 1 hour daily, either for 1 day (PO 1d) or for 30 consecutive days (PO 30d). Control mice were handled similarly with water exposure. (B) Novelty-suppressed feeding (NSF) test: Following the exposure paradigm, 24hr later, mice were placed in a novel open-field arena with a food pellet located in the center. Latency to begin feeding was measured as latency to feed (s). (C) Elevated plus maze (EPM) test: avoidance behavior was assessed by measuring time spent in open and closed arms, and total distance traveled. Data are shown as mean ± SEM, n=10-26. Statistical significance determined by one-way ANOVA followed by Dunnett’s multiple comparisons test, *p<0.05, ***p<0.01, ***p<0.001.

### Activity-dependent mapping of predator odor responsive ensembles in whole mouse brain

Predator odor (PO) is primarily detected through the vomeronasal system, a specialized chemosensory pathway that processes social and threat-related olfactory cues (19,20). The vomeronasal organ (VNO) transmits information to the accessory olfactory bulb, which in turn projects to limbic structures involved in defensive behaviors (19,20). To identify the neural ensembles activated by predator odor exposure, we performed whole-brain Fos mapping (Fig.2) using immunostaining of serial slices and analysis following a single 1-hour exposure to predator odor (PO) or water (Ctl). Several brain regions (Supplementary Table 1) exhibited significant increases in Fos+ neurons compared to controls in acute predator odor samples (Fig.2A-B), including the insular cortex (IC), lateral septum (LS), cingulate cortex (Cing), and paraventricular hypothalamus (PVH). Fold change analysis (Fig. 2C) of the number of Fos+ neurons in each brain region shows that, among these regions, the LS and PVH had the largest differences in Fos expression with a substantial increase relative to controls (>40 fold change; p=0.04 for LS and PVH). Moderate to high increases in Fos were also significant for other brain regions, including the cingulate cortex (Cg), parabrachial nucleus (PBN), basolateral amygdala (BLA), lateral hypothalamic area (LHA), suprammamillary nucleus (SUM), ventral tegmental area (VTA), paraventricular nucleus of the thalamus (PVT), pontine nuclei (Pnt), insular cortex (IC), lateral entorhinal cortex (LEC), raphe pallidus nucleus (RPa), medial preoptic area (MPO), periaqueductal gray (PAG) and superior colliculus (SC). These findings highlight the neural activation triggered by predator odor exposure and suggest a prominent role for the limbic system and its downstream targets in processing predator-related stress.

**Figure 2.**
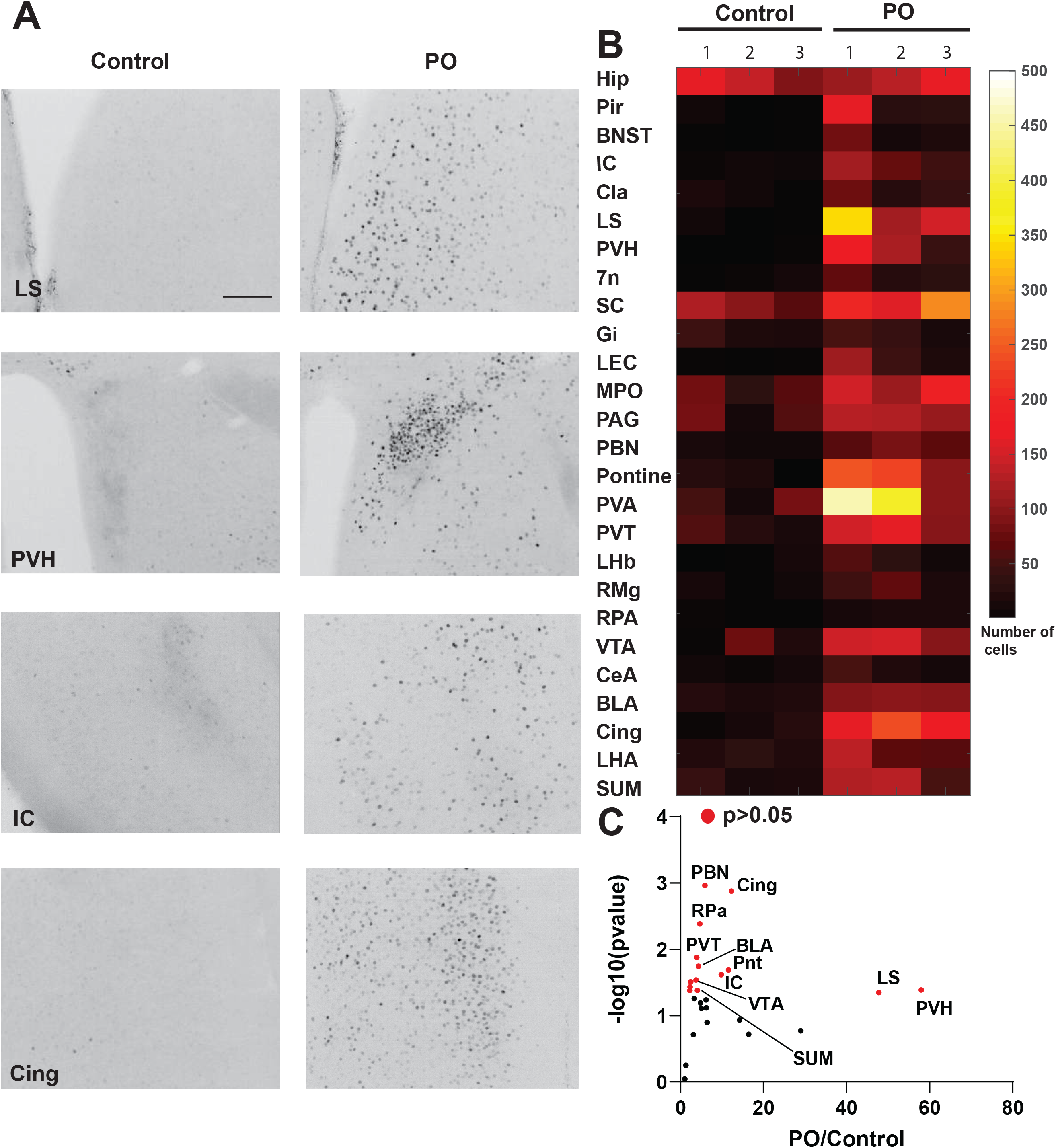
Predator odor exposure induces activation of several brain regions involved in stress response and other behaviors. (A) Representative coronal sections showing Fos immunoreactivity in selected brain regions of control (Ctl) and predator odor-exposed (PO) mice. Exposure to predator odor markedly increased the number of Fos-positive nuclei in multiple forebrain and midbrain regions including the piriform cortex (Pir), insular cortex (IC), lateral septum (LS), bed nucleus of the stria terminalis (BNST), and paraventricular hypothalamic nucleus (PVH). Scale bar = 150 µm. (B) Heatmap of Fos+ cell counts across multiple brain regions in control and PO-exposed mice (n = 3 per group). Regions are organized along the y-axis and individual biological replicates along the x-axis. Color scale represents number of Fos+ cells counted, from low (black) to high (yellow). (C) Volcano plot summarizing the fold change (x-axis) and significance (–log10 p-value, y-axis) of Fos expression across all analyzed regions. P-value was calculated using unpaired Student’s t-test (control vs PO). Red points denote significantly activated regions (p<0.05).

### Identification of predator odor-responsive neuronal clusters in the LS using activity-dependent and single-cell nuclei RNAseq

Based on our whole brain Fos mapping data and our previous work showing that LS neurons integrate stress signals to suppress feeding (23), we reasoned that the LS would be the prime candidate for integrating and relaying predator odor information downstream to control avoidance behaviors, in the presence or absence of food. Thus, we decided to further dissect the molecular identity of predator odor-responsive neurons in the lateral septum (LS) using PhosphoTRAP, a technique that enables the immunoprecipitation of translating ribosomes from activated neuronal ensembles (23,30). PhosphoTRAP selectively isolates mRNA from neurons that are actively engaged in signaling pathways involving phosphorylated ribosomal protein S6 (pS6), a marker of neurons undergoing synaptic transmission (30). This approach allows for the targeted capture of molecular signatures specific to neurons activated, in this case, by predator odor exposure (Fig.3A). First, to ensure that LS neurons exhibit increased pS6 levels upon predator odor exposure, we performed immunohistochemistry against pS6 after a single 1-hour exposure to predator odor and observed a significant increase in pS6 levels in the LS when compared to controls (Fig. 3A; **p=0.0017). This is consistent with Fos data indicating increased neuronal activation in response to predator odor. We next performed bulk RNA sequencing on LS tissue dissected from mice exposed to a single 1-hour predator odor session, or water-exposed controls. Transcriptomic analysis identified an enrichment of activity-dependent genes (Fig.3B), including Fos and Egr4, as well as genes encoding neuropeptides and neurotransmitters. Among these we observed enrichment in genes encoding for neuropeptides such as neurotensin (Nts) and corticotrophin-releasing hormone (Crh) in PO-exposed samples versus controls (Fig.3B). Neurotensin ranks among the top neuropeptides that have increased mRNA levels in PO-exposed immunoprecipitation (IP) samples versus input (INP), when compared to control IP versus INP samples (Fig.3B and C). We also analyzed gene markers that were enriched in PO-exposed samples that characterize specific LS populations (Fig.3C) Our analysis shows that PO-activated LS neurons express several neuropeptides such as Nts, Crh, glucagon-like peptide receptor 1 (Glp1r), Crh receptor 2 (Crhr2), somatostatin (Sst) and cocaine and amphetamine regulated transcript (Cartpt). We analyzed another recently characterized LS marker, mu-opioid receptor 1 (Oprm1; 31), but this specific LS marker was depleted in PO-exposed samples. Although these neuropeptide genes are enriched in PO-exposed samples, they, with the exception of Crh and Nts, are not significantly enriched when PO-exposed samples are compared to controls (Fig. 3B). Among the classes of genes that encode for neurotransmitters, GABA is enriched in our PO-activated samples compared to controls while Glutamate (Glu), Glycine (Gly), serotonin (5ht) and dopamine (DA) are depleted (Fig.3C). These results corroborate previous research showing the involvement of GABAergic LS neurons that co-express these neuropeptides in other modalities of stress (23–27,31).

**Figure 3.**
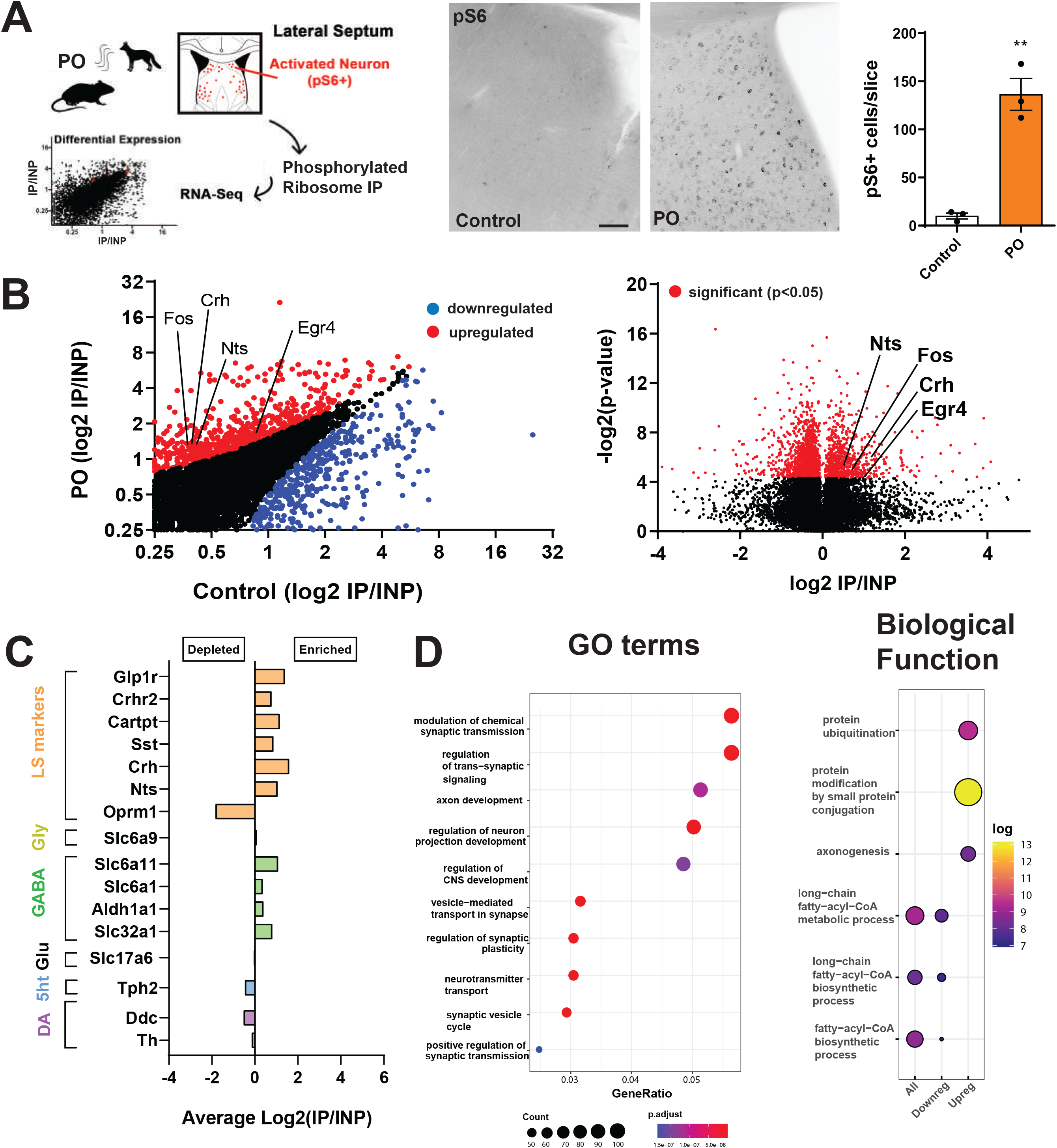
PhosphoTRAP analysis reveals predator odor-induced transcriptional activation of stress-and neuromodulatory genes in the lateral septum. (A) Experimental design and validation of neuronal activation. Mice were exposed to predator odor (PO) for 1 hour, followed by collection of the lateral septum (LS) for PhosphoTRAP (pS6-immunoprecipitation of ribosome-bound mRNAs). Left: schematic showing pS6+ neuron isolation and RNA sequencing workflow. Middle: representative immunohistochemical images showing increased phosphorylated S6 (pS6)-positive neurons in the LS following PO exposure compared to controls. Right: quantification of pS6+ cell per slice. Scale bar = 100 µm. Data are shown as mean ± SEM, n=3. Unpaired Student’s t-test, **p<0.01. (B) Transcriptomic profiling of activated LS neurons. Left: scatter plot of normalized gene expression (log₂ IP/INP) for control versus PO conditions. Red dots show upregulated transcripts and blue dots represent downregulated transcripts (p<0.05) Right: volcano plot showing significant differential expression of genes enriched in PO-activated LS neurons (red, p<0.05). Specific transcripts are shown including *Fos, Crh, Nts,* and *Egr4*. (C) Neurochemical signature of activated LS neurons is shown as average log₂(IP/INP) ratios highlighting genes enriched (right) or depleted (left) in the PO-activated LS population. (D) Functional enrichment analysis of PhosphoTRAP datasets. Left: Gene Ontology (GO) analysis. Right: biological function enrichment.

To gain deeper insight into the functional pathways altered by predator odor exposure, we conducted Gene Ontology (GO) analysis of the top 100 PO-enriched genes (Fig. 3D). This revealed an enrichment of genes involved in the modulation of synaptic transmission and axonogenesis, suggesting adaptive changes in neuronal communication within the LS following exposure to predator odor. Among biological processes that were upregulated by predator odor (Fig. 3E), protein modification by small-protein conjugation and protein ubiquitination are among the top processes upregulated in the LS by 1h predator odor exposure. These categories cover SUMO-, NEDD8-and ubiquitin-ligase components, suggesting that predator odor rapidly engages proteostatic mechanisms that tag synaptic proteins for turnover or trafficking which is important for synapses and neuropeptide release (32). Conversely, we observed a downregulation of genes associated with metabolic processes, indicating a potential shift in cellular energy utilization, such as long-chain fatty acid synthesis and metabolism. This rapid shift in energy utilization is also evidenced by upregulation of genes that enhance catabolism and substrate-shuttling within the cell, thus predicting a metabolic pivot toward other types of energetic substrates, such as glucose. Predator odor significantly enriched for genes (Supplementary Fig. 1) involving cellular metabolism such as Free Fatty Acid Receptor 3 (Ffar3), Gamma-Glutamyltransferase 1 (Ggt1), Adenosylmethionine Decarboxylase 1 (Amd1), Solute Carrier Family 38 Member 11 (Slc38a11), Carnosine Dipeptidase 1 (Cndp1), Vitamin D Receptor (Vdr), G Protein-Coupled Receptor 35 (Gpr35). Several of these genes are directly involved in substrate utilization and redox balance. Ffar3, a receptor for gut-derived short-chain fatty acids, was elevated, suggesting enhanced LS sensitivity to peripheral metabolic signals during stress (33). Ggt1, which regulates glutathione cycling, was strongly upregulated, consistent with an increased demand for antioxidant defenses under conditions of high neuronal activity (34). Cndp1, encoding carnosine dipeptidase, also increased, pointing to altered histidine and β-alanine metabolism, which could support oxidative stress mitigation (35). Amd1, a key enzyme in polyamine synthesis, also showed upregulation, suggesting an upregulation in the requirement for polyamines in synaptic plasticity and structural remodeling(36,37). Nutrient transport pathways were affected as well. Slc38a11, a sodium-coupled amino acid transporter, was elevated, suggesting enhanced availability of amino acid substrates for neurotransmitter synthesis during predator-induced activation (38). Finally, two regulatory genes with broader signaling roles, Vdr and Gpr35, were upregulated. Vdr contributes to calcium signaling and gene regulation, and its increase may promote neuroprotection and synaptic stability in the LS (39). Gpr35, a receptor for kynurenine pathway metabolites and bile acids, may link stress-induced metabolic changes with neuromodulatory control of LS circuits(40). Most of these genes are reported to be expressed in non-neuronal cells such as glia, endothelial cells but also neurons, suggesting a rapid metabolic change in cells that support neuronal function and in neurons themselves. Altogether, these findings suggest that predator odor exposure triggers a rapid LS transcriptional program that balances synaptic remodeling with metabolic adaptation.

Next, to comprehensively determine which specific LS population was recruited during predator odor exposure, we used snRNA seq data generated by the Allen Brain Institute (ABC Atlas;41) and cross referenced their dataset with our PhosphoTRAP dataset (Fig.4A). Similar to what has been described by other groups (31,25), our snRNA seq data re-analysis identified discrete cell types in the septal complex defined by their unique gene expression enrichment. Based on our PhosphoTRAP data, our analysis focused on GABAergic subtypes, and we identified 6 molecularly defined neuronal subtypes in the LS that are GABAergic. Among these neurons, we found within the Slc32a1 cluster (GABA+), cell types that corresponded to previously identified LS populations (Fig. 4A-D; 31,24,25), including neurotensin+ (Nts+;Slit2 cluster), glucagon-like peptide 1 receptor+ (Glp1r+; Zeb2 cluster), mu-opioid receptor 1+ (Oprm1+;Otx2), corticotropin-releasing hormone receptor 2 (Crhr2+; Nkx2-1), Met proto-oncogene (Met+; Lmo1 cluster), paired box 6+ (Pax6+, Pax6 cluster) neurons (Fig.4C and D). Among these specific populations, other genes were identified, at different proportions and expression levels, as possible markers for each one of these populations such as neuromedin B (Nmb), dachshund 2 (Dach2), calcitonin receptor (Calcr), melanocortin receptor 3 (Mc3r) and galanin (Gal) (Fig.4B). We also observed overlap between many clusters (Fig.4B-D) such as Sst+, Glp1r+ and Nts+ clusters. This overlap can also be observed in spatial transcriptomics analysis from MERFISH experiments using wild-type C57BL6/J mice and imputed gene data (data from ABC Atlas, 42, Supplementary Fig.2).

**Figure 4.**
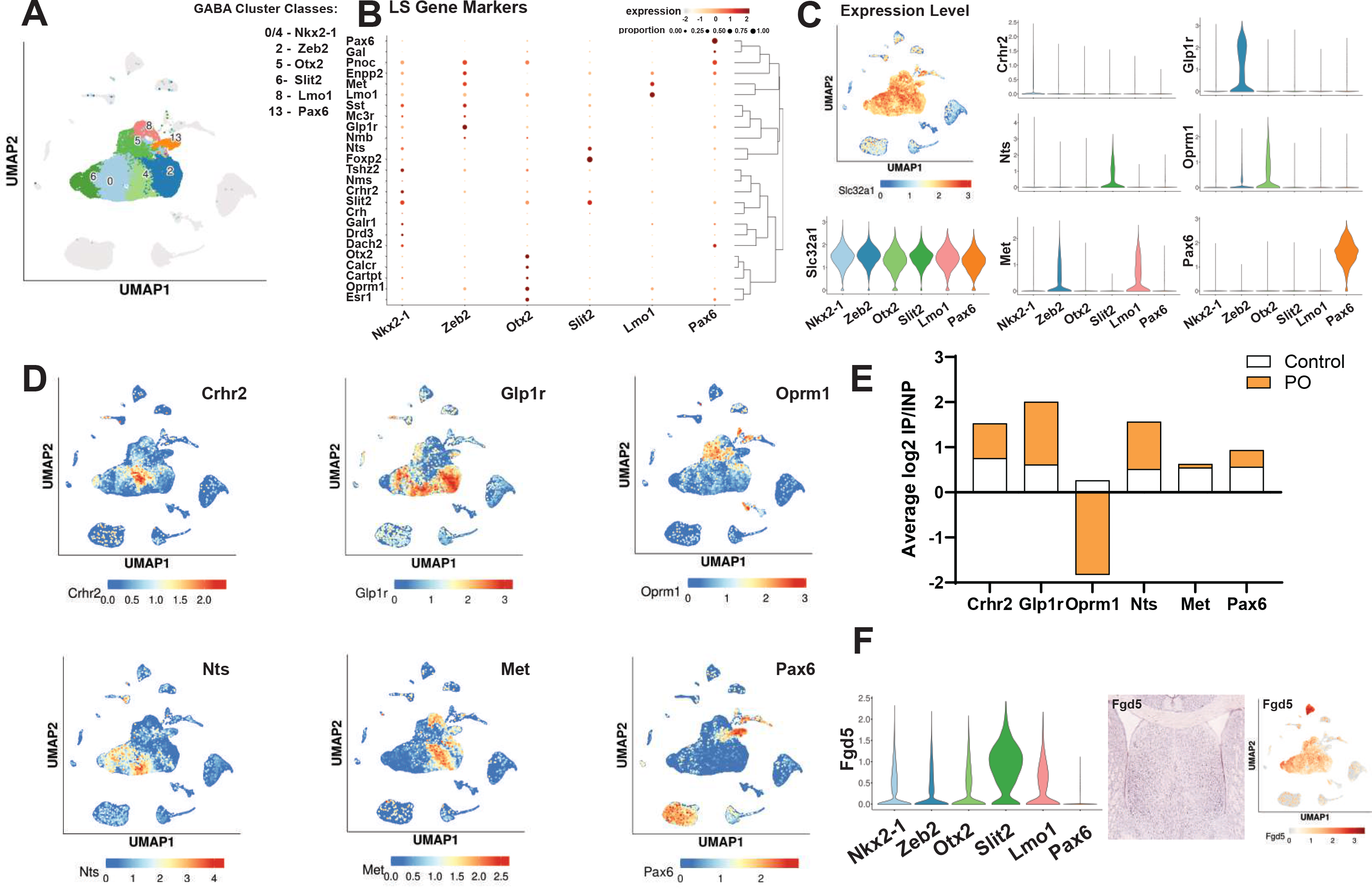
Single-nuclei RNA-seq analysis reveals molecular identity of stress-activated subtypes of lateral septum (LS) neurons. (A) UMAP of LS GABAergic neuron clusters. Reanalysis of publicly available single-nuclei RNA-seq data from the Allen Brain identified multiple transcriptionally distinct GABAergic LS subpopulations (clusters 0-13). Major cluster classes were annotated by transcription factor expression, including *Nkx2-1*, *Zeb2*, *Otx2*, *Slit2*, *Lmo1,* and *Pax6*. (B) Dot plot of LS cluster gene markers shows expression levels (color intensity) and proportion of expressing cells (dot size) for LS marker genes, including neuropeptides (*Crh, Nts, Gal, Sst*), receptors (*Glp1r, Crhr2, Oprm1*), and transcriptional regulators (*Pax6*). (C) Cluster-specific gene expression patterns are shown in violin plots to depict the expression of *Slc32a1* (the vesicular GABA transporter) across LS clusters. Expression patterns of genes (*Crhr2, Glp1r, Nts, Oprm1, Met, Pax6*) reveal subtype-specific enrichment within clusters. (D) Spatial distribution of neuromodulatory receptors and peptides are shown through UMAP highlighting the cluster-specific localization of *Crhr2, Glp1r, Oprm1, Nts, Met,* and *Pax6* transcripts. (E) Integration with PhosphoTRAP data with single-nuclei RNA seq data show the averaged log2(IP/INP) ratios from PhosphoTRAP sequencing (Control vs PO) demonstrating selective enrichment of transcripts throughout the LS clusters. (F) Integration from PhsphoTRAP data and single nuclei RNA seq data identified a novel LS Slit2 marker (Fgd5) and in situ hybridization image from Allen Brain ISH Database show strong *Fgd5* expression within LS. Violin plot of *Fgd5* expression confirms this gene as a marker for *Slit2*-defined LS clusters.

We next performed a cross-referencing analysis of the LS marker genes identified in the snRNA-seq analysis with our PhosphoTRAP datasets from PO-responsive neurons to identify cluster markers that were predominantly enriched in our datasets (Fig.4E). Among these specific populations, genes significantly enriched in PO-exposed samples were found to be enriched in the Crh2r+, Glp1r+ and Nts+ clusters, while PO-exposed genes were less enriched in the Met+ and Pax6+ clusters and genes enriched in PO-exposed samples were not found in the Oprm1+ cluster (Fig.4E). Using this analysis, we found that predator odor exposure recruited mainly 3 clusters (Nkx2-1, Zeb2 and Slit2 corresponding to Crh2r+, Glp1r+ and Nts+ clusters) in the LS, while not recruiting, or even inhibiting, the Otx2 (corresponding to Oprm1+) LS cluster. Among these specific genes, novel transcripts were found, such as Fgd5 (FYVE, RhoGEF and PH domain-containing 5; Fig.4F). Fgd5 is highly enriched in Slit2+ clusters but can also be found with lower expression levels in other LS clusters with exception of Pax6+ clusters. Fgd5 encodes a protein that primarily activates CDC42 (Cell division control protein 42 homolog), probably driving cytoskeleton remodeling in neurons (43).

Based on our data (Fig. 3 and 4) and previous work from us and others (23–25) showing that Nts+ populations co-express many of the molecules enriched in PO-exposed samples and integrate stress signals, we selected LS^NT^ neurons as the prime candidate population for encoding predator odor information in the LS and modulating avoidance behaviors.

### Neurotensin-expressing LS (LS^NT^) neurons are necessary and sufficient for chronic predator-induced avoidance in mice

Next, we decided to orthogonally validate the functional role of LS^NT^ neurons in chronic predator odor-induced avoidance. First, to evaluate whether LS^NT^ neurons were activated by predator odor, we exposed mice to predator odor or control (water) acutely and performed in situ hybridization for Fos and NT (Fig.5A-C). We found that ∼40 cells/slice of NT+ neurons in the LS also expressed Fos after 1h of predator odor exposure, which represent about one third of the whole NT+ population in the LS (Fig.5C, **p=0.0031). Next, using fiber photometry in Nts-cre mice injected in the LS with a cre-dependent AAV encoding GCaMP6s (Fig.5D) we found that LS^NT^ neurons reliably show a large calcium signal in response to predator odor exposure during a 5 min presentation protocol (*p=0.04), but not in response to water (control). This increase in activity was rapidly present in the first seconds of odor presentation. These results suggest that LS^NT^ neurons are robustly and rapidly activated by predator odor.

**Figure 5.**
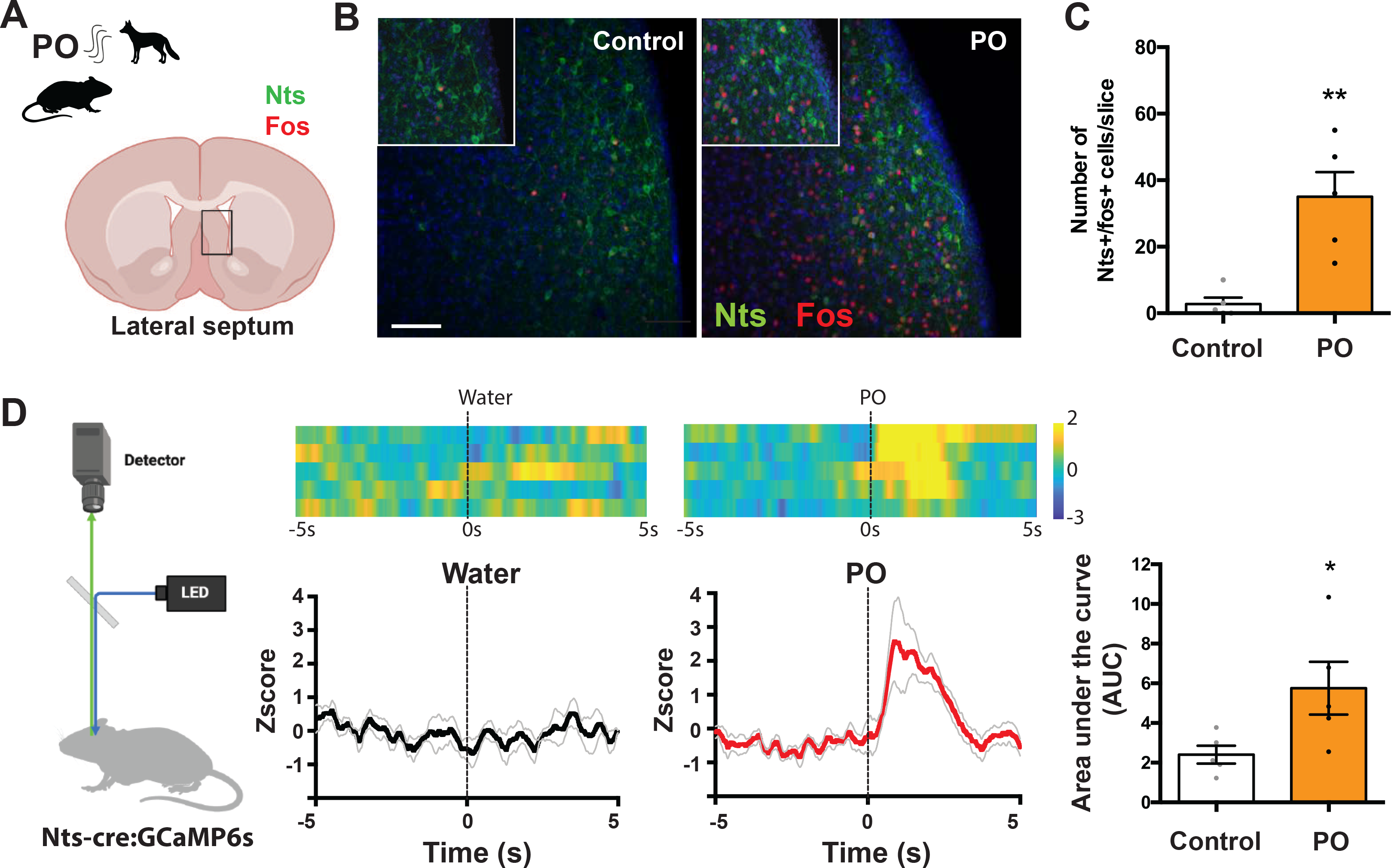
Predator odor activates Nts-expressing neurons in the lateral septum and drives rapid calcium responses in vivo. (A) Schematic of the experimental design. Mice were exposed to predator odor (PO) or control conditions, and activation of Nts-expressing neurons in the lateral septum (LS) was assessed using Fos immunostaining. (B) Representative images of LS sections showing Nts⁺ neurons (green), Fos⁺ nuclei (red), and DAPI (blue) in control (left) and PO-exposed (right) mice. Inset show higher-magnification view of Nts/Fos co-labeling. Scale bar, 100 µm. (C) Quantification of Nts⁺/Fos⁺ double-positive neurons, data are shown as mean ± SEM, Students t-test, n=4/group, **p < 0.01. (D) Fiber photometry recordings in Nts-Cre:GCaMP6s mice during control (water) and PO exposure. Top: heatmaps showing single-trial z-scored fluorescence aligned to stimulus onset (time 0). Bottom: average zscore traces (± SEM). Right: area under the curve (AUC) analysis, data are shown as mean ± SEM, Students t-test, n=5/group, *p < 0.05.

To test whether activation of LS^NT^ neurons is sufficient to mimic the behavioral effects of chronic predator odor exposure, we injected Nts-cre mice in the LS with cre-dependent AAV encoding for an activatory DREADD (hM3Dq), or mCherry as a control, to selectively activate LS^NT^ neurons. Mice were then treated with C21 (1mg/kg i.p.) chronically for 30 days. As controls, we showed that C21 (1 mg/kg i.p.) was sufficient to significantly activate Nts+ neurons in the LS (Fig. 6A-C; **p=0.0019). Following this activation protocol, we performed NSF and EPM tasks, 24hrs after the last C21 injection, to evaluate avoidance behaviors in injected mice in the absence of predator odor. We found that chronic, daily activation of LS^NT^ neurons increased the latency to feed in the NSF task similarly to predator odor (*p=0.04, Fig,6D) and decreased the time spent in the open arms of EPM (*p=0.04), without changing time in closed arms (p=0.49) or locomotion (p=0.83, Fig. 6E). These data suggest that chronic LS^NT^ neuron stimulation alone is sufficient to induce the same avoidance behaviors as chronic exposure to predator odor and in the absence of predator odor.

**Figure 6.**
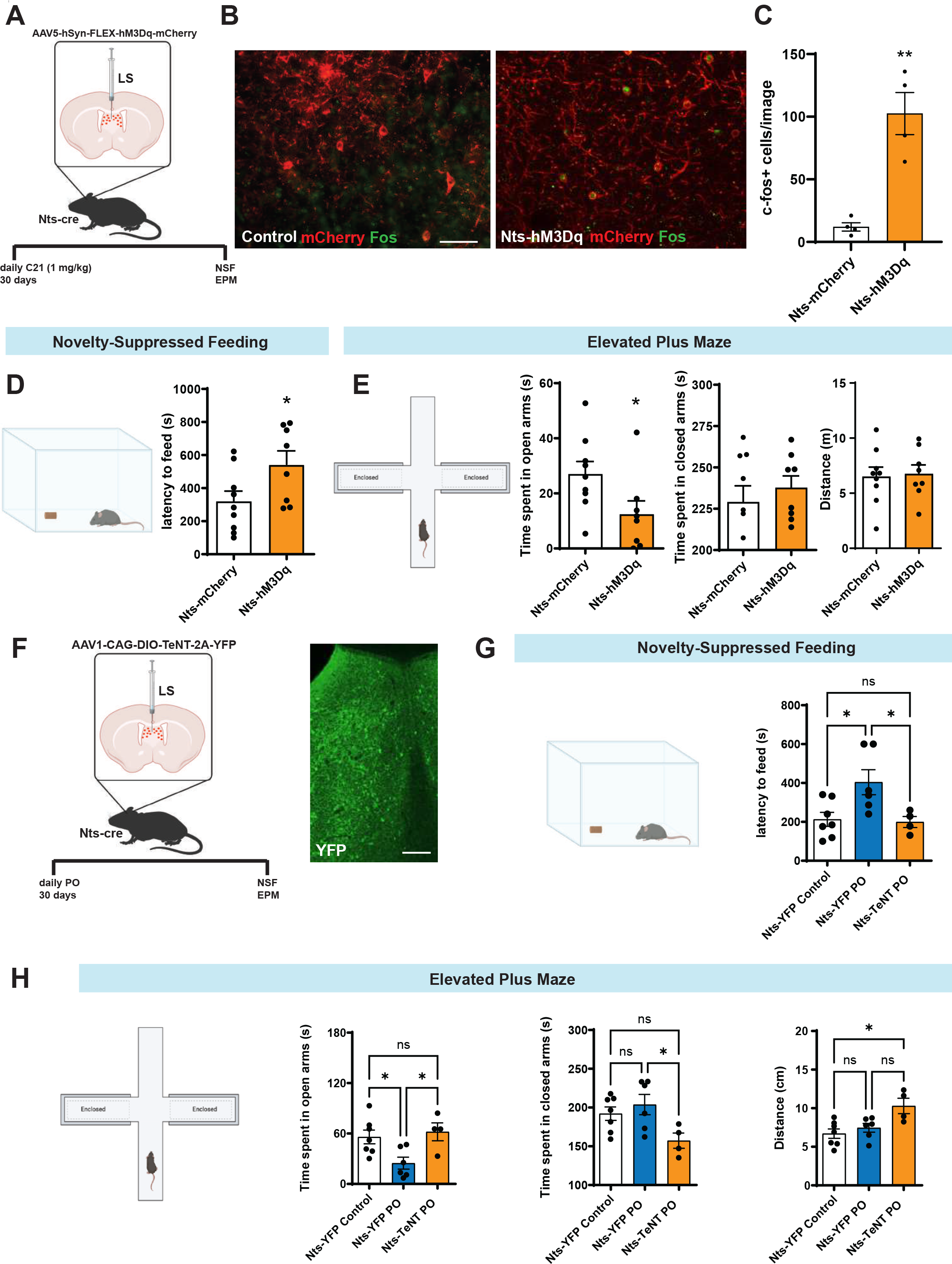
Nts-expressing LS neurons regulate avoidance behaviors. (A) Experimental design for chemogenetic activation of LS Nts neurons. Nts-Cre mice received bilateral LS injections of AAV5-hSyn-FLEX-hM3Dq-mCherry or control virus followed by chronic C21 administration (1 mg/kg/day, i.p.). Behavioral testing was performed 30 days later. (B) Representative LS sections showing mCherry-labeled neurons (red) and Fos immunoreactivity (green) in control and hM3Dq-expressing mice. Chemogenetic activation robustly increases Fos expression in LS Nts neurons. Scale bar, 50 µm. (C) Quantification of Fos⁺/mCherry⁺ cells, data are shown as mean ± SEM, Unpaired students t-test, n=4, **p< 0.01. (D) Novelty-suppressed feeding test, data are shown as mean ± SEM, Unpaired students t-test, n=8-9, *p <0.05. (E) Elevated Plus Maze (EPM) performance (open arms, closed arms, distance), data are shown as mean ± SEM, Unpaired students t-test, n=8-9, *p < 0.05. (F) Experimental design for synaptic silencing of LS NT neurons using TeNT concomitant with predator odor (PO) exposure. Nts-Cre mice received bilateral LS injections of AAV1-CaMKII-DIO-TeNT-2A-GFP or control virus. Right: GFP expression in LS confirming viral spread. Scale bar, 100 µm. (G) Novelty-suppressed feeding after PO exposure, data are shown as mean ± SEM, One-way ANOVA with Bonferroni post hoc test, n=4-7, *p<0.05). (H) Elevated Plus Maze after PO exposure (open arms, closed arms, distance), data are shown as mean ± SEM, One-way ANOVA with Bonferroni post hoc test, n=4-7, *p < 0.05.

Next, we silenced LS^NT^ neurons synaptically to determine whether they are necessary for the behavioral effects of chronic PO exposure. To chronically inhibit LS^NT^ neurons, we injected Nts-cre mice in the LS bilaterally with a cre-dependent AAV encoding for tetanus toxin (TeNT; 44), to irreversibly silence LS^NT^ neuron’s synaptic activity (Fig.6F). Control animals were injected with AAVs encoding for YFP alone. After recovery from surgery, TeNT or YFP mice were exposed to 30 days of predator odor for 1 hour per day. A third group injected into the LS with YFP alone was exposed to water for 30 days as control. Following this protocol, we performed NSF and EPM tasks to evaluate avoidance behaviors in injected mice (Fig.6F). We found that silencing LS^NT^ neurons during chronic daily exposure to predator odor prevented the development of avoidance behaviors exhibited in control animals (Fig.6G, Nts-YFP Control vs Nts-TeNT PO, p=0.99 and Nts-YFP PO vs Nts-TeNT PO, *p=0.04). In the EPM task, we observed similar preventative effects of inhibiting LS^NT^ neurons. Silencing LS^NT^ neurons (Nts-TeNT PO) prevented the behavioral effects of chronic predator odor, resulting in increased time spent in the open arms (Fig.6H, Nts-YFP Control vs Nts-TeNT PO, p=0.99 and Nts-YFP PO vs Nts-TeNT PO, *p=0.03) while decreasing time in closed arms of the EPM task (Fig.6H, Nts-YFP Control vs Nts-TeNT PO, p=0.15 and Nts-YFP PO vs Nts-TeNT PO, *p=0.04), and increasing locomotion (Fig.6H, Nts-YFP Control vs Nts-TeNT PO, *p=0.01 and Nts-YFP PO vs Nts-TeNT PO, p=0.0512). These data suggest that LS^NT^ neurons are necessary to mediate development of avoidance behaviors induced by chronic predator odor exposure.

### Downstream mapping of functionally relevant predator odor-responsive LS^NT^ neurons

To identify the downstream circuits that receive inputs from PO-responsive neurons and control behavioral output, we used TRAP2 mice (45) injected in the LS bilaterally with a cre-dependent AAV carrying an axon-filling mCherry to indelibly tag predator odor-activated neurons and their axonal projections (Fig.7A-D), irrespective of molecular identity. Predator odor-responsive cells in the LS were indelibly labeled with mCherry upon 4-OHT injection and exposure to predator odor for 6 hours. Control animals were injected with 4-OHT and exposed to water for 6 hrs (Fig.7A). We showed that predator odor exposure labeled a dense number of neurons in the LS (average 100-200 cells across the whole LS extension compared to controls, *p=0.04), reinforcing our transcriptomics data suggesting that different LS sub-populations respond to predator odor exposure. Using whole brain mCherry mapping and serial images, we found that mCherry+ neurons in the LS robustly projected to the infralimbic cortex (IL), the dorsal hippocampus region (dHip), the lateral hypothalamus (LH), the periaqueductal gray (PAG) and the suprammamillary nucleus (SUM) of mice (Fig.7B-D). On average, the densest projections were observed in the lateral hypothalamus (LH) and the suprammamillary nucleus (SUM) suggesting that these two hypothalamic regions may be potential candidates in mediating the effects of predator odor on behavior. Additionally, our previous work and others suggest that both LH and SUM together with PAG are known targets for LS^NT^ neurons, while IL and dHip are targets for other LS populations such as Sst+ LS neurons (46). These data also corroborate our transcriptomics data that suggest that the LS region has a diversity of cell types with specific outputs and diverse functions (23–25,31).

**Figure 7.**
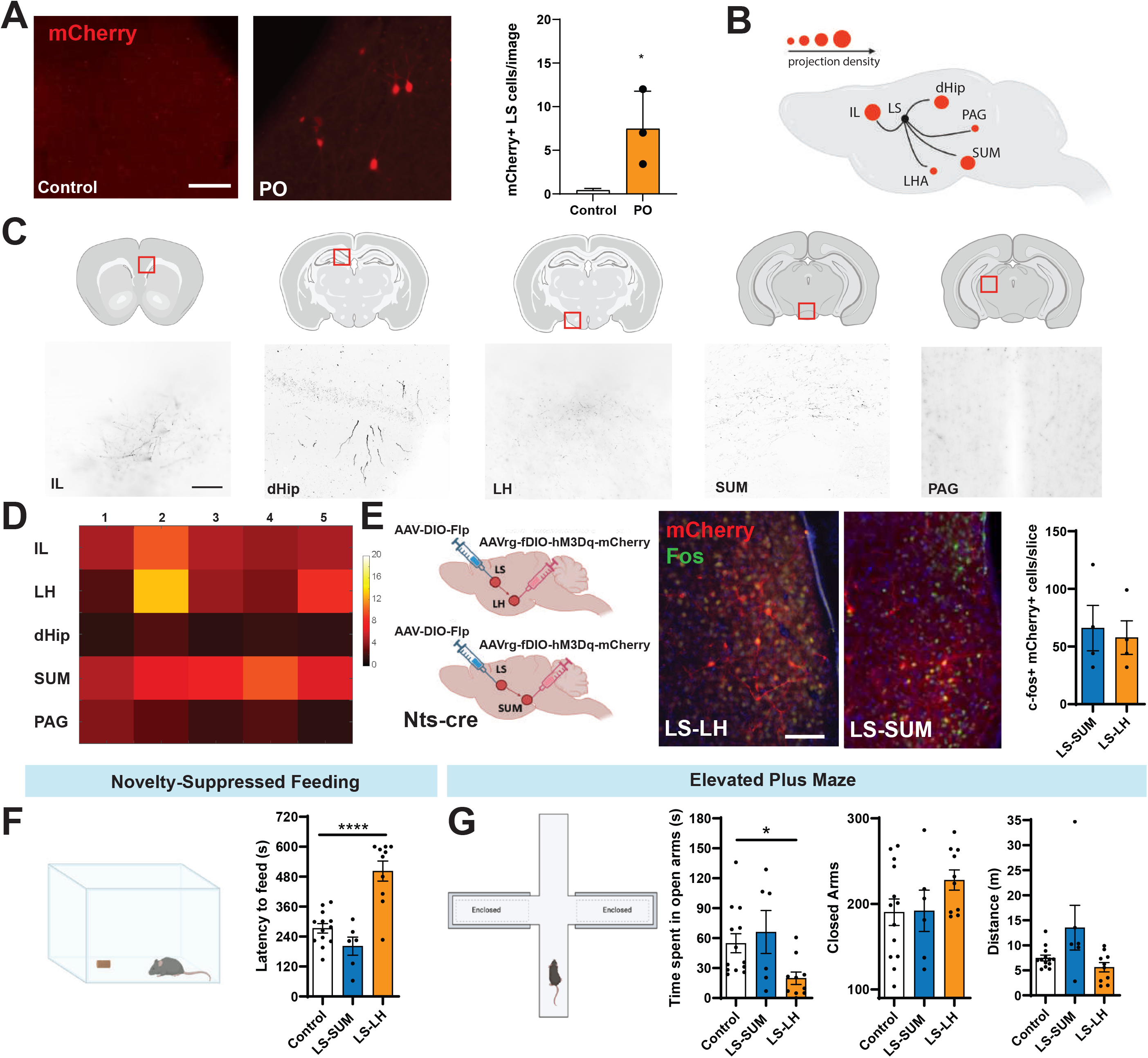
Predator odor-responsive LS neurons project to stress-and anxiety-related regions and drive behavioral avoidance through LS→LH, but not LS→SUM pathways. (A) Activity-dependent tagging of predator odor-responsive LS neurons using TRAP2 mice. TRAP2 mice were bilaterally injected in the LS with AAV5-hSyn-DIO-mCherry, exposed to PO or water, and mCherry expression was induced to label recently activated LS cells. Left: representative LS images. Right: quantification of mCherry⁺ neurons per slice, data are shown as mean ± SEM, Unpaired Students t-test, n=3, *p < 0.05. Scale bar, 50 µm. (B) Schematic depicting major projection targets of PO-activated LS neurons identified through whole-brain axon mapping, including infralimbic cortex (IL), dorsal hippocampus (dHip), lateral hypothalamus (LH), supramammillary nucleus (SUM), and periaqueductal gray (PAG). (C) Representative brain sections showing mCherry-labeled LS axons within identified projection sites (IL, dHip, LH, SUM, PAG). Insets (red boxes) highlight corresponding anatomical regions. Scale bar, 50 µm. (D) Heatmap summarizing projection density across all mapped targets. Brighter colors indicate stronger innervation, revealing dominant projections to LH and SUM. (E) Pathway-specific activation of LS projections. Left: intersectional viral strategy using AAV-DIO-Flp bilaterally injected into the LS of Nts-cre mice and AAVrg-fDIO-hM3Dq-mCherry injected into LH or SUM to express hM3Dq selectively in LS NT+ neurons projecting to each target. Middle: representative images of LS NT+ neurons expressing hM3Dq-mCherry following retrograde viral delivery. Right: quantification of Fos⁺/mCherry⁺ cells after chemogenetic activation confirms engagement of LS→LH and LS→SUM neurons (n=4). Scale bar, 100 µm. (F) Novelty-suppressed feeding, data are shown as mean ± SEM, One-way ANOVA with Bonferroni post hoc test, n=6-13, ****p<0.0001), indicating enhanced anxiety-like avoidance. (G) Elevated Plus Maze (open arms, closed arms, distance), data are shown as mean ± SEM, One-way ANOVA with Bonferroni post hoc test, n=6-13, *p < 0.05.

To specifically target LS^NT^ neurons, which we previously identified as being an important LS population integrating predator odor signals and inducing avoidance, projecting to the LH or SUM we used an intersectional approach consisting of a bilateral LS injection in Nts-cre mice of AAVs encoding cre-dependent Flp and an injection in the SUM or LH of a retrograde AAV encoding Flp-dependent activatory DREADDs (Fig.7E-G). Using this strategy, we observed that we could consistently target a similar number of LS^NT^-projecting neurons both to LS and SUM (average 50 cells per 50 microns of brain slice, Fig.7E). After surgery recovery, we used chronic, daily C21 (1mg/kg, i.p.) injections for 30 days to evaluate which specific LS^NT^ projection was relevant for avoidance behaviors, in a similar manner to chronic exposure to predator odor in mice. Following this activation protocol in LS^NT^-SUM or LS^NT^-LH cohorts, we observed that chronic activation of LS^NT^-LH circuit, but not LS^NT^-SUM, mimicked the behavioral performance observed in the NSF (Fig.7F, Control vs LS-SUM, p=0.42; Control vs LS-LH, ****p=0.0001) and EPM tasks similarly to experiments chronically activating cell body LS^NT^ neurons and chronic predator odor exposure for 30 days (Fig.7G, open arms, Control vs LS-SUM, p=0.99; Control vs LS-LH, *p=0.04; closed arms, Control vs LS-SUM, p=0.99; Control vs LS-LH, p=0.16, distance, Control vs LS-SUM, p=0.08; Control vs LS-LH, p=0.99). In fact, chronic activation of LS^NT^-SUM circuit trended towards anxiolytic behaviors in the NSF and EPM tasks, but without reaching statistical significance (Fig.7F and G). These results suggest that LS^NT^ neurons integrate aversive information from chronic predator odor exposure and relay it downstream to LH neurons to modulate avoidance behaviors in mice.

### LS neurotensin is required for predator-odor induced avoidance in mice

Next, we asked whether neurotensin itself could be an important molecular mechanism driving these changes in behavior in addition to being a useful molecular marker of neuron populations mediating exploratory behavior and stress response. LS^NT^ neurons are both neurotensinergic and GABAergic (23), thus the possibility of two different molecular mechanisms is conceivable. Additionally, data analysis mined from the Allen Brain Atlas for ISH (47) suggests that both the LH and SUM contain NTSR1 receptors, with the SUM containing the highest density of NTSR1 (Supplementary Fig. 3). Thus, we decided to evaluate whether neurotensin itself was required for the effects observed in mice chronically exposed to predator odor. To this end, adult Nts-flox/flox mice received bilateral stereotaxic injections of an AAV-hSyn-Cre-GFP in the LS (Fig.8A). Cre-mediated recombination excised the floxed Nts exon, selectively eliminating neurotensin in LS neurons (LS^NT^KO). Littermates injected with the same volume of AAV-hSyn-Cre-GFP served as controls and were only exposed to water for 30 days. Three weeks later, cohorts were exposed chronically to daily predator odor or water for 30 days. Twenty-four hours after the final exposure, mice were tested on the EPM and then, NSF tasks to assess avoidance behaviors.

**Figure 8.**
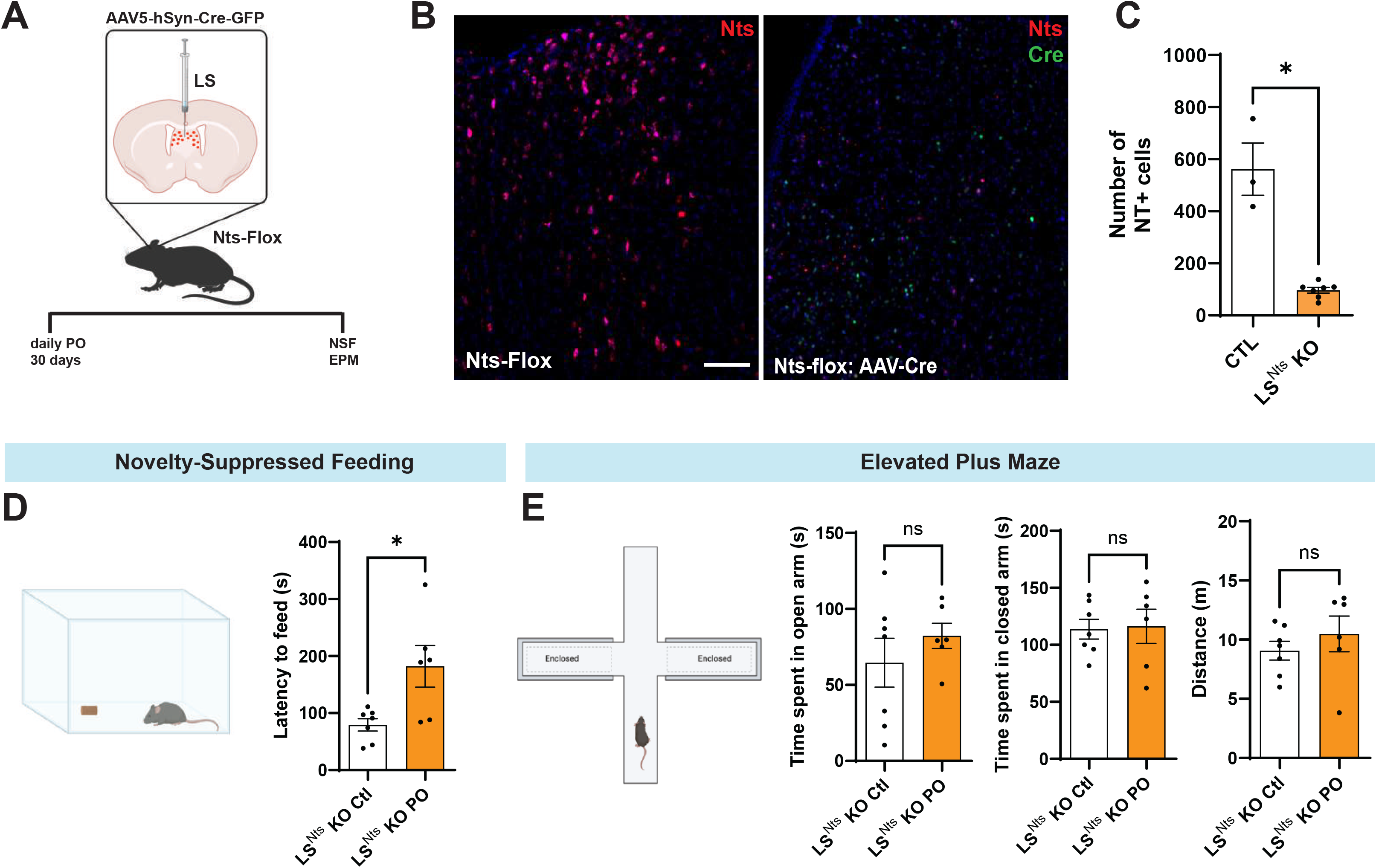
Neurotensin is required to regulate chronic PO exposure-induced avoidance. (A) Experimental design for conditional deletion of neurotensin (Nts) in the lateral septum (LS). Nts-flox/flox mice received bilateral LS injections of AAV5-hSyn-Cre-GFP, followed by 30 days of daily predator odor (PO) exposure prior to behavioral testing (novelty-suppressed feeding, NSF; elevated plus maze, EPM). (B) Representative LS images showing Nts immunostaining (red) or GFP (green) in control Nts-flox/flox mice (left) and near-complete loss of Nts signal following LS-targeted Cre expression (right). Scale bar, 100 µm. (C) Quantification of Nts⁺ cells in LS^Nts^ KO and control mice, data are shown as mean ± SEM, Unpaired students t-test, n=3-7, *p<0.05. (D) Novelty-suppressed feeding test, data are shown as mean ± SEM, Unpaired students t-test, n=6-7, *p < 0.05. (E) Elevated plus maze performance (open arms, closed arms, distance), data are shown as mean ± SEM, Unpaired students t-test, n=6-7, ns, p>0.05.

We found that chronic predator odor significantly increased the latency to begin feeding in the NSF task (Fig.1B, latency to feed = ∼400s) compared with unstressed controls (Fig.1B, latency to feed = ∼200s). In LS^NT^ KO exposed to 30d of PO, however, we observed the same latency to feed (Fig.8D, latency to feed = ∼200s) as the unstressed mice (Fig.1B) and were even significantly lower in LS^NT^ KO unstressed animals (Fig.8D, latency to feed = ∼100s, p<0.05). We then asked whether the same manipulation would influence avoidance in the EPM (Fig.1C; time in open arms =∼30 sec average; time in closed arms =∼190 sec average; distance = ∼14 m average), and this shift was completely prevented when neurotensin was deleted from the LS (Fig.8D and E, LS^NT^ KO PO and Ctl; time in open arms =∼90 sec average; time in closed arms =∼110 sec average; distance = ∼10 m average). Thus, our data suggests that neurotensin released by LS neurons is required for avoidance behaviors in the EPM and NSF tasks induced by chronic predator odor exposure.

These data identify LS neurotensin signaling as a critical driver of the chronic stress-induced avoidance, thus leading to a reduction in exploration in novel environments.

## DISCUSSION

Threat detection occurs constantly, with animals weighing the prospective value of exploration of novel environments against the possibility of danger(3). Here we identify a genetically defined, LS neurotensinergic (LS^NT^) circuit that imposes a behavioral “brake” on exploration and increases avoidance when danger is persistent, and we delineate the molecular, cellular and projection mechanism by which this “brake” is engaged.

### A cell-specific circuit for avoidance under persistent threat

Using predator odor as an ethologically relevant stressor in mice (19,20), we found that many brain regions are engaged in predator odor response (Fig.2) and that behavior drifts from exploration to avoidance after 30 consecutive days of exposure (Fig.1). Among the brain regions, the LS was one of the regions with the highest Fos induction differences observed. Activity-dependent translating ribosome capture (PhosphoTRAP) and snRNA-seq (Fig.3 and 4) revealed that predator odor activates several LS populations, but it enriches for transcripts that are concentrated mostly in GABAergic, Nts+ ensembles. Our transcriptomic data also revealed that not only LS^NT^ neurons are activated by predator odor in the LS, clusters that are marked by the expression of Sst (expressed in Zeb2, Nkx2-1, Slit2 and Otx2 clusters), Glp1r (Zeb2 cluster), Cartpt (Nkx2-1 and Otx2 cluster) and Crh (Nkx2-1 cluster) are also activated. Our previous work using in situ hybridization (23) confirms the snRNA seq data, suggesting that within the LS, many of these clusters overlap to some extent with Nts+ populations (23). Interestingly, Oprm1+ neurons (Otx2 cluster) seem to be not enriched or even inhibited by predator odor exposure, suggesting that possibly this neuronal population may be either part of a local inhibitory circuit or receives inhibitory input from upstream regions activated by predator odor. Despite this, it remains unknown whether these clusters form a local inhibitory network, self-regulating their own activity and responses to sensory or interoceptive stimuli.

Evidence from our data suggests a causal role of LS^NT^ neurons in threat avoidance induced by predator odor. First, in vivo calcium photometry demonstrates robust activation of LS^NT^ neurons whenever mice encounter predator odor (Fig.5). Second, chemogenetic excitation of these neurons for 30 consecutive days is sufficient to replicate the avoidance phenotype normally observed after a 30-day predator odor exposure (Fig.6). Third, chronic synaptic silencing of LS^NT^ neurons during predator odor exposure completely blocks the transition from exploration to avoidance (Fig.6). Fourth, selective deletion of NT in LS neurons prevents predator-induced avoidance (Fig.8). Together, these results strongly suggest that, although other LS populations may also be activated by predator odor, LS^NT^ neurons are both necessary and sufficient to mediate persistent threat avoidance after chronic exposure to predator odor or chemogenetic activation.

Using anatomical tracing using TRAP2 mice to indelibly tag predator odor-responsive LS neurons and intersectional DREADD manipulations we further identified a direct LS^NT^◊LH projection as the critical downstream pathway. Sustained activation of this monosynaptic pathway alone reproduces the behavioral effects of chronic predator exposure and global LS^NT^ neuronal stimulation (Fig.7). Interestingly, LS^NT^◊SUM projections showed a neutral or even an opposite trend towards anxiolytic behavior, without reaching significance. Both projection sites are rich in NTSR1 (Supplementary Fig.3), which further points to the functional diversity of NT projections and NTSR1 circuits in the brain. These findings suggest a specific neurotensinergic circuit that suppresses exploratory drive, increasing avoidance, via targeted inhibition of LH targets. However, the identity of LH and SUM neurons that receive neurotensinergic projections are still unknown and whether these downstream effects depend on NTSR1 are currently being investigated.

### Novel insights into the stress-induced acute remodeling of septal circuits

Threat detection forces the brain to reconcile two competing demands: it must reshape synaptic networks quickly enough to bias behavior away from danger, yet it must perform these changes using finite metabolic reserves(48). Our transcriptomic analysis of the LS following a 1h predator odor exposure suggests a possible mechanistic framework for how these metabolic demands may be met, acutely. Using bioinformatics to analyze Gene Ontology and Biological Process enrichment, we showed a possible mechanism by which molecular adaptations following a stressor may occur (Fig.3). The biological function of up-regulated transcripts is almost exclusively for protein posttranslational modifications and axonogenesis, suggesting increased proteostatic turnover and cytoskeletal remodeling that can modify or remodel synaptic strength or morphology within the first hours of threat exposure. In contrast, downregulated genes map onto long-chain fatty-acyl-CoA biosynthesis and metabolism pathways, indicating an immediate decrease in lipid metabolism. The suppression of lipid biosynthetic genes points to an unexpected metabolic mechanism that may drive septal function. By transiently reducing lipid synthesis pathways, LS neurons may preserve its ATP and NADPH supplies to engage in more metabolic remanding tasks such as oxidative stress control (which demands NADPH) and synaptic function (which can generate reactive oxygen species), illustrating how metabolic and synaptic homeostasis are co-regulated during acute threat (48,49). These results are corroborated with an increase in genes that are associated with synaptic transmission, but remain to be validated experimentally.

These results may therefore delineate a parallel metabolic response in LS neurons upon threat exposure: (1) fast synaptic rewiring driven by enhanced post-translational modification and neurite-growth pathways, and (2) metabolic re-prioritization that diverts resources away from NADPH-expensive lipid synthesis toward synaptic function. The convergence of these cellular and systems findings implies that sensory cues, such as predator odor, may trigger a rapid remodeling of septal circuits, transforming the LS function from goal-motivated exploration into arrest and avoidance behavior over time. Whether these pathways are important for avoidance or synaptic remodeling remains unknown. However, these data provide molecular and metabolic insights into how behaviorally important circuits are rapidly remodeled when activated by sensory stressors in an acute manner. Future work will assess the effects of chronic PO exposure in circuit remodeling combining in vivo proteomics with detailed synaptic analysis will be essential to determine whether the transcriptional signature we observe translates into the predicted reorganization of LS synapses and whether similar programs are engaged by other sensory threat modalities.

### Neurotensin as a modulatory switch

Our data suggests that GABAergic, NT-containing neurons are important for maladaptive behaviors following a prolonged threat signal. High-affinity NTSR1 and NTSR2 receptors are abundantly expressed in LH neurons that govern arousal and foraging (50,51, Supplementary Fig.3), providing an anatomically privileged target for septal NT. Moreover, peptide release could be triggered by burst firing and sustained presynaptic Ca²⁺ elevation(52), both conditions that may occur and would characterize LS activity during predator odor exposure, thus it remains to be tested. In addition, NT shows a marked Ca²⁺ dependence in brain preparations (53). Thus, together, these properties make the GABA/NT axis a suitable candidate for chemically communicating stressful information downstream to LH and other target regions. Indeed, we show here that deleting NT in LS neurons prevented avoidance after chronic PO exposure (Fig. 8), implying that the neuropeptide NT, and perhaps not GABA per se, engages LH circuits during chronic predator odor exposure. However, it remains to be seen whether deletion of vGAT in LS^NT^ neurons would have any effect in predator odor-induced avoidance behaviors. One possible model, which remains to be investigated, is the synaptic mechanisms by which NT co-released with GABA (54) shifts the synaptic threshold that separates exploration and avoidance behavioral outputs, especially in scenarios that switch from acute to chronic. This is important because in our previous work (23) we did not see an effect of acutely activating LS NT+ neurons in EPM tasks, suggesting that synaptic plasticity events may be needed to shift behavior from exploration to avoidance. Based on these, targeting the neurotensinergic signaling axis may therefore represent a rational strategy for modulating maladaptive avoidance without broadly dampening limbic excitability.

### Limitations and future directions

In the present study, we cannot yet rule out contributions from LS co-peptides (Crh, Sst and Cartpt) or from GABA itself to avoidance behaviors mediated by LS^NT^ neurons. Determining whether neurotensin acts pre-or post-synaptically in LH, whether NTSR1 or NTSR2 plays a role in LS^NT^-LH circuits remains unknown. In addition, understanding how different stress modalities (restraint, forced swim, noise, fasting) engage LS neurons to modulate avoidance would be beneficial to elucidate a common circuit and molecular mechanisms that shift behavior from exploration to avoidance, during acute or chronic exposure scenarios. This would be essential for targeting and developing new therapies to treat stress-related disorders. Finally, sex differences in predator-stress responses were not addressed in our work, thus, given higher anxiety prevalence in females (55), mapping LS^NT^ dynamics across sexes would be an interesting next step.

## Conclusion

Our work positions LS^NT^-LH circuits and septal NT as a molecular and circuit mechanism through which persistent threat from predator odor suppresses exploration of novel environments. By specifying the cell type, transmitter and downstream target that convert chronic stress into avoidance, we provide a framework for mechanistic interrogation of maladaptive coping and open avenues for targeted intervention in stress-related pathology.

## STAR*METHODS

Detailed methods are provided and include the following:

- KEY RESOURCES TABLE
- CONTACT FOR REAGENT AND RESOURCE SHARING
- EXPERIMENTAL MODEL AND SUBJECT DETAILS
- METHOD DETAILS

- 2,4,5-trimethylthiazole exposure
- Behavioral Tasks: Novelty-suppressed feeding and Elevated plus maze
- Immunohistochemistry and Image quantification
- PhosphoTRAP
- SnRNA-seq reanalysis
- In vivo fiber photometry
- Chemogenetics
- Cell Tagging
- Stereotaxic injections and Optic Fiber Implantation
- Functional Annotation of RNA Sequencing Data
- QUANTIFICATION AND STATISTICAL ANALYSIS
- DATA AND SOFTWARE AVAILABILITY

- Data Resources

## STAR∗METHODS

### KEY RESOURCES TABLE

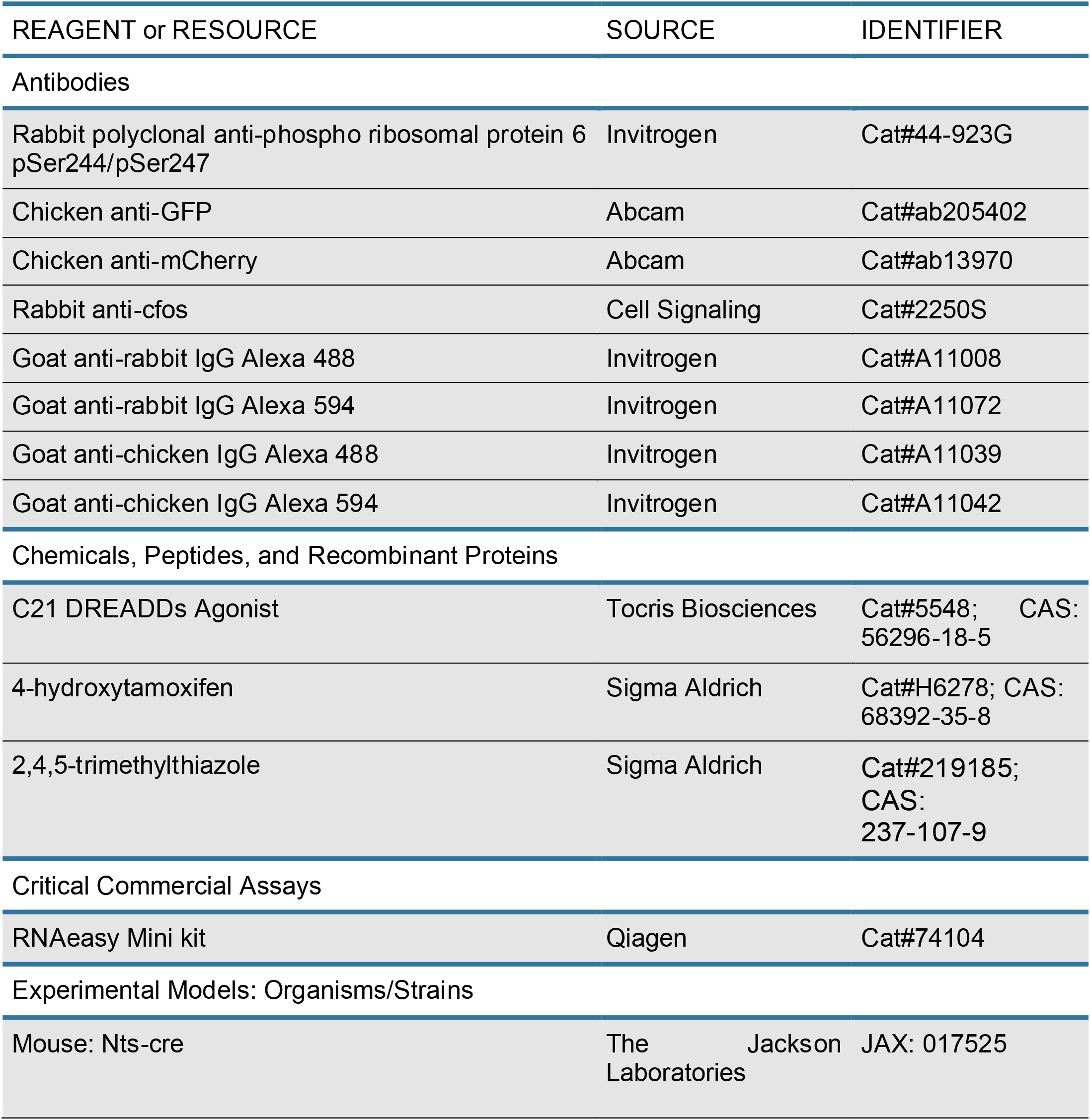
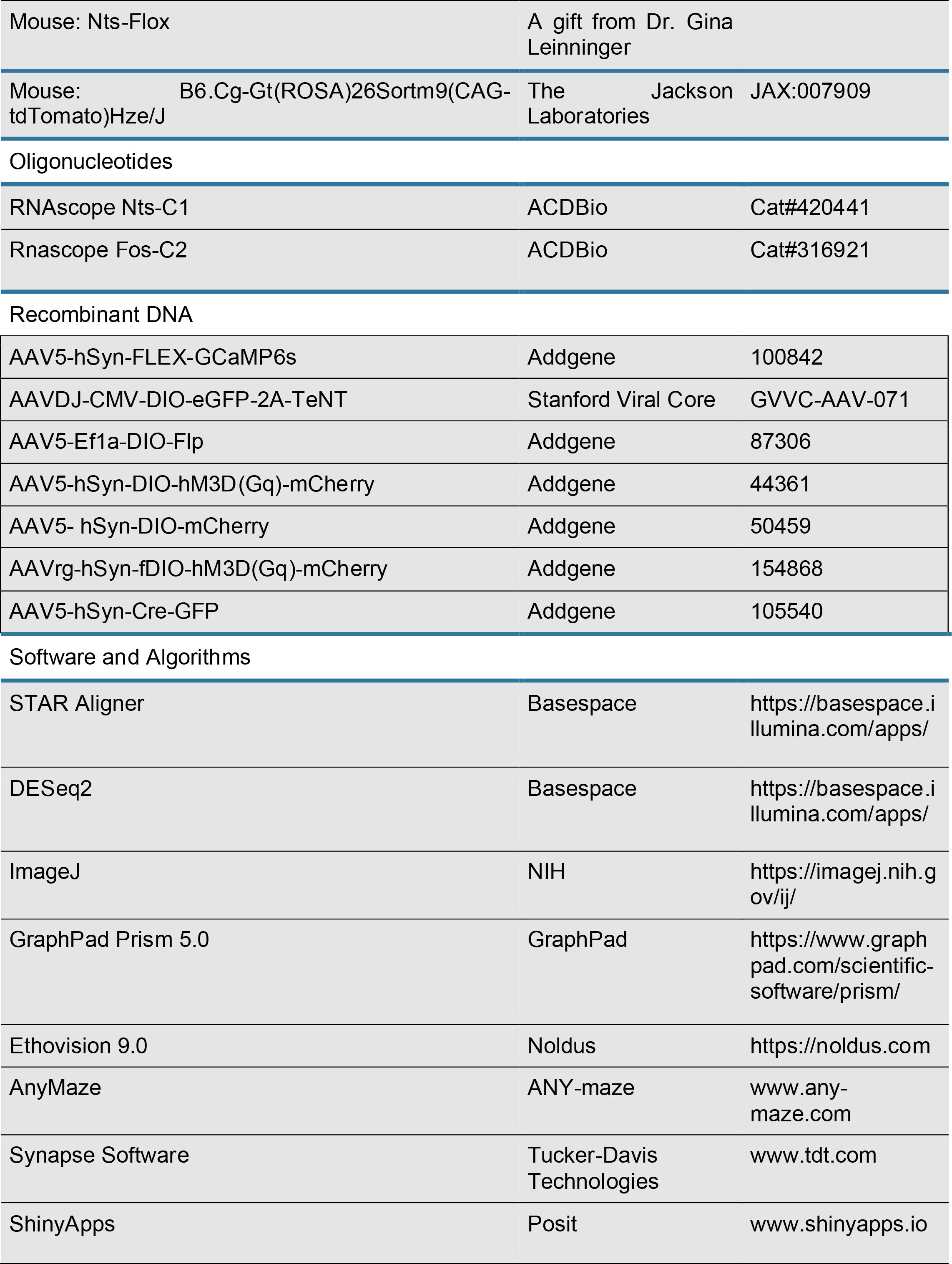
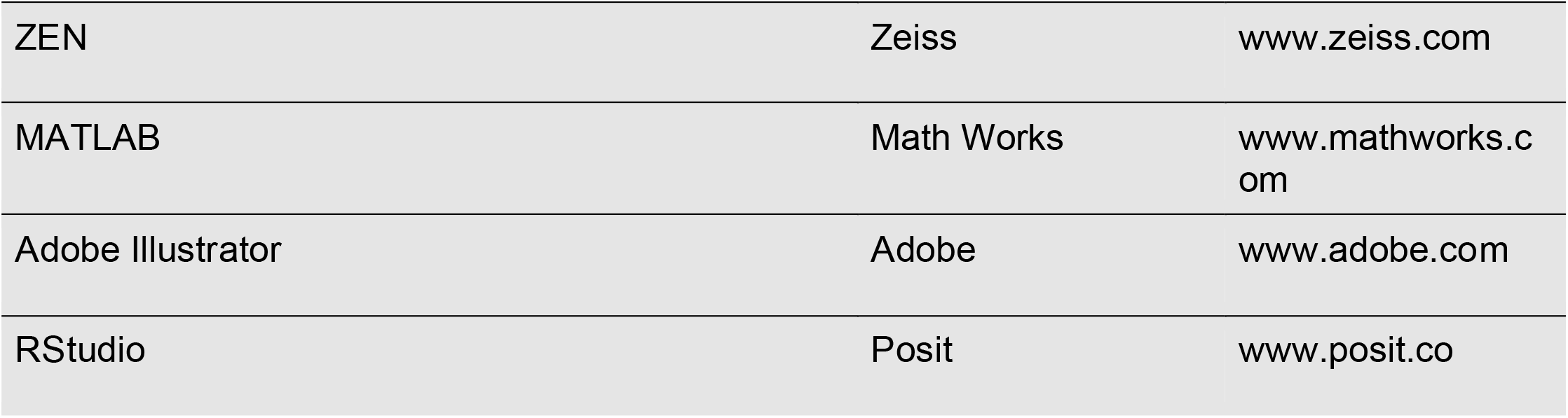

| REAGENT or RESOURCE | SOURCE | IDENTIFIER |
| --- | --- | --- |
| Antibodies |  |  |
| Rabbit polyclonal anti-phospho ribosomal protein 6 pSer244/pSer247 | Invitrogen | Cat#44-923G |
| Chicken anti-GFP | Abcam | Cat#ab205402 |
| Chicken anti-mCherry | Abcam | Cat#ab13970 |
| Rabbit anti-cfos | Cell Signaling | Cat#2250S |
| Goat anti-rabbit IgG Alexa 488 | Invitrogen | Cat#A11008 |
| Goat anti-rabbit IgG Alexa 594 | Invitrogen | Cat#A11072 |
| Goat anti-chicken IgG Alexa 488 | Invitrogen | Cat#A11039 |
| Goat anti-chicken IgG Alexa 594 | Invitrogen | Cat#A11042 |
| Chemicals, Peptides, and Recombinant Proteins |  |  |
| C21 DREADDs Agonist | Tocris Biosciences | Cat#5548; CAS: 56296-18-5 |
| 4-hydroxytamoxifen | Sigma Aldrich | Cat#H6278; CAS: 68392-35-8 |
| 2,4,5-trimethylthiazole | Sigma Aldrich | Cat#219185; CAS: 237-107-9 |
| Critical Commercial Assays |  |  |
| RNAeasy Mini kit | Qiagen | Cat#74104 |
| Experimental Models: Organisms/Strains |  |  |
| Mouse: Nts-cre | The Jackson Laboratories | JAX: 017525 |
| Mouse: Nts-Flox | A gift from Dr. Gina Leininger |  |
| Mouse: B6.Cg-Gt(ROSA)26Sortm9(CAG-tdTomato)Hze/J | The Jackson Laboratories | JAX:007909 |
| Oligonucleotides |  |  |
| RNAscope Nts-C1 | ACDBio | Cat#420441 |
| Rnascope Fos-C2 | ACDBio | Cat#316921 |
| Recombinant DNA |  |  |
| AAV5-hSyn-FLEX-GCaMP6s | Addgene | 100842 |
| AAVDJ-CMV-DIO-eGFP-2A-TeNT | Stanford Viral Core | GVVC-AAV-071 |
| AAV5-Ef1a-DIO-Flp | Addgene | 87306 |
| AAV5-hSyn-DIO-hM3D(Gq)-mCherry | Addgene | 44361 |
| AAV5- hSyn-DIO-mCherry | Addgene | 50459 |
| AAVrg-hSyn-fDIO-hM3D(Gq)-mCherry | Addgene | 154868 |
| AAV5-hSyn-Cre-GFP | Addgene | 105540 |
| Software and Algorithms |  |  |
| STAR Aligner | Basespace | https://basespace.illumina.com/apps/ |
| DESeq2 | Basespace | https://basespace.illumina.com/apps/ |
| ImageJ | NIH | https://imagej.nih.gov/ij/ |
| GraphPad Prism 5.0 | GraphPad | https://www.graphpad.com/scientific-software/prism/ |
| Ethovision 9.0 | Noldus | https://noldus.com |
| AnyMaze | ANY-maze | www.any-maze.com |
| Synapse Software | Tucker-Davis Technologies | www.tdt.com |
| ShinyApps | Posit | www.shinyapps.io |
| ZEN | Zeiss | www.zeiss.com |
| MATLAB | Math Works | www.mathworks.com |
| Adobe Illustrator | Adobe | www.adobe.com |
| RStudio | Posit | www.posit.co |

## CONTACT FOR REAGENT AND RESOURCE SHARING

Further information and requests for reagents may be directed to, and will be fulfilled by the corresponding author Estefania Azevedo.

## EXPERIMENTAL MODEL AND SUBJECT DETAILS

Mice were housed according to the guidelines of MUSC’s Animal Facility (DLAR), in a 12 hr light/dark cycle and with free access to standard chow diet and water. C57BL/6J background males and females at age 10-14 weeks were used throughout this study and all animals experiments were approved by the MUSC IACUC following the National Institutes of Health guidelines for the Care and Use of Laboratory Animals. For behavioral studies only male mice were used. C57BL/6J wild-types (Cat#000664), Nts-cre (51 Stock# 017525), Fos2A-iCreER (TRAP2; Cat# 030323) were purchased from Jackson Laboratories (Bar Harbor, ME). Nts-Flox/Flox was a gift from Dr. Gina Leinninger and is currently available at JAX Stock# 036262.

## METHODS DETAILS

### 2,4,5-trimethylthiazole exposure

2,4,5-trimethylthiazole (predator odor, PO) is a synthetic chemical analog of the red fox-derived aversive odor 2,4,5-trimethylthiazoline (TMT) that produces stress and innate fear responses (14–16). Similar to TMT, PO produces robust responses that are easily quantified and controlled when compared to natural predator odors, greatly reducing problems with variability encountered with natural predator stimuli. Mice were exposed to undiluted (3uL) PO in a filter paper in an air-tight cage under a chemical hood to avoid odor dispersion for 1h acutely or 1h/day for 30 days. Control mice were exposed in the same context to water for equivalent times. After PO exposure, mice were transferred to their home cage and housed accordingly.

### Behavioral Tasks: Novelty-suppressed feeding and Elevated plus maze

For novelty-suppressed feeding (NSF) task, mice were fasted for 16h and then placed in an acrylic box with two pellets of standard chow. Latency to approach and consume the food was recorded using an overhead camera for 10 minutes and manually scored by a blinded experimenter. For elevated plus maze (EPM), mice were placed at the center of a cross-shaped, elevated maze in which two arms are closed with dark walls and two arms are open and allowed to explore for 5 min. Afterward, the mice were returned to their home cage and the maze floor was cleaned in between subjects. All subjects were tested by a blinded experimenter and monitored using a camera and behavior (time spent in open and closed arms, distance, and velocity) were analyzed using ANY-maze (Stoelting). Latency to approach food, time spent in the open arms and closed arms and distance traveled were plotted using GraphPad Prism 9.0 (GraphPad).

### Immunohistochemistry and Image quantification

Mice were perfused, and brains were postfixed for 24 hr in 10% formalin. Brain slices were taken using a vibratome (Leica), blocked for 1 hr with 0.3% Triton X-100, 3% bovine serum albumin (BSA), and 2% normal goat serum (NGS) and incubated in primary antibodies for 24 hr at 4°C in a shaker. Then, serial free-floating slices were washed three times for 10 min in 0.1% Triton X-100 in PBS (PBS-T), incubated for 1 hr at room temperature with secondary antibodies, washed in PBS-T and mounted in Vectamount with DAPI (Southern Biotech). Antibodies used here were: anti-c-fos (1:500; Cell Signaling), anti-GFP (1:1000, Abcam), anti-mCherry (1:1000; Abcam) goat-anti-rabbit (Alexa 488 or Alexa 594, 1:1000) or goat anti-chicken Alexa488 or Alexa594 (1:1000). Images were taken using AxioImager.M2 microscope (Zeiss) and images were processed using ImageJ software (NIH, Schneider et al., 2012). C-fos counts were conducted using ImageJ and validated by a blind experimenter in whole brain in serial slices per animal (min 3 mice/group) and areas annotated according to Allen Brain Atlas. For axonal quantification, using threshold masking segmented axons were quantified and mean grey value/slice was calculated for each registered brain region in a minimum of 5 mice/group to account for potential variability in injection site locations or viral expression across samples.

### PhosphoTRAP

PhosphoTrap (Activity-based transcriptomics) profiling experiments were performed accordingly (23,30). Briefly, mice were separated into groups of 6 mice per group, exposed to PO for 1 hr and sacrificed immediately following the stress. Naive mice were exposed to water in the same context as PO animals and sacrificed after 1h. After euthanasia, brains were removed, and septal area were dissected on ice and pooled (6 brain samples per experiment replicate, total of 12 brains). Tissue was homogenized and clarified by centrifugation. Ribosomes were immunoprecipitated using 4 mg of polyclonal antibodies against pS6 (Invitrogen, #44-923G) previously conjugated to Protein A-coated magnetic beads (Thermofisher). A small amount of tissue RNA was saved before the immunoprecipitation (Input) and both input and immunoprecipitated RNA (IP) were then purified using RNAeasy Mini kit (QIAGEN) and RNA quality was checked using a RNA PicoChip on a bioanalyzer. RIN values >7 were used. cDNA was amplified using SMARTer Ultralow Input RNA for Illumina Sequencing Kit and sequenced on an Illumina NextSeq platform. RNA sequencing raw data was uploaded and analyzed using STAR aligner and DESeq2 in BaseSpace (Illumina) using an alignment to annotated mRNAs in the mouse genome (UCSC, *Mus musculus* assembly mm10). The average immunoprecipitated (IP) and Input value of each enriched and depleted genes with a q-value higher than 0.05 were plotted using GraphPad Prism 9.0 (GraphPad). Neurons identified using PhosphoTRAP were further validated using in situ hybridization, and functional testing.

### snRNA-Seq reanalysis

Lateral septum snRNA Seq data was reanalyzed from Allen Brain Map published data (41). Raw Striatum (WMB-10Xv3-STR-raw.h5ad) dataset containing lateral septum tissue from Whole Mouse Brain (WMB) 10X version 3 (10Xv3) single-cell RNA sequencing (scRNA-seq) was downloaded. Downstream analysis was performed in R with Seurat (v4.3.0) (56) using customized R scripts. Seurat object was made from these raw counts, subset of LSX region was used for this analysis, metadata from cell_metadata_with_cluster_annotation.csv was added to LSX subset by matching cell barcode. Ensemble ID was converted to gene name for convenience and seurat object was filtered for protein coding genes only. Quality Filtering was conducted by retaining cells that had unique molecular identifiers (UMIs) less than 60000, number of features / Genes (nGene) greater than 250 and mitochondrial content less than 10 percent. Doublets were removed using scDblFinder (v1.12.0) (57) for a resultant expression matrix with 21,477 genes and 50,842 cells. The SCTransform workflow was used for count normalization, initial integration, and to identify highly variable genes (58) using 30 principal components and resolution of 0.5 for Louvain clustering and UMAP. Cluster marker genes were identified using FindAllMarkers using Wilcoxon Rank Sum test with the standard parameters. We identified 21 final distinct clusters, 6 LS GABA clusters and cell annotation from the original metadata was used to label the clusters. Cell type deconvolution of BulkRNA expression data using this scRNA as reference was performed using R package MuSiC (v1.0.0) (59).

### In vivo fiber photometry

Mice were acclimated to patch cables for 5 min for 3d before experiments. Analysis of the signal was done using the fiber photometry system and processor (RZ5P) from TDT (Tucker-Davis Technologies), which includes the Synapse software (TDT). The bulk fluorescent signals from each channel were normalized to compare across animals and experimental sessions (23). The 405 channel was used as the control channel. GCaMP6s signals that are recorded at this wavelength are not calcium-dependent, thus, changes in signal can be attributed to autofluorescence, bleaching, and fiber bending. Accordingly, any fluctuations that occurred in the 405 control channels were removed from the 465 channel before analysis. Change in fluorescence (DF) was calculated as (465 nm signal – fitted 405 nm signal), adjusted so that a DF/F was calculated by dividing each point in DF by the 405-nm curve at that time. Z-scores were calculated as (signal – signal median)/ median absolute deviation to account for variability across animals. Behavioral variables were timestamped in the signaling traces via the real-time processor as TTL signals from Noldus (Ethovision) software. This allowed for precise determinations of the temporal profile of signals in relation to specific behaviors. Behaviors were tested as follows. First, we tested the response to water presented in a filter paper for 5 min in a clean home cage. We then tested PO presentation in a filter paper for 5 min. All behaviors were manually scored by a second investigator who did not conduct the photometry experiment in a blinded manner. Calcium signals and behavioral data were time-locked and peri-events calculated for each animal. Codes for data extraction and analysis were performed using custom codes in MATLAB.

### Chemogenetics

For chronic chemogenetics experiments, C21 DREADDS Agonist (Tocris Biosciences) in saline at 1 mg/kg dose i.p. was used per mice. Injections were given daily in the afternoon for 30 days. Control animals (reporter only) received the same dose and number of injections as subjects. At day 31 and 32, animals were tested in the EPM and NSF tests, respectively. C21 efficiency was validated after experiments using anti-c-Fos immunostaining.

### Cell Tagging

AAV5-hSyn-DIO-hM3Dq–mCherry (Addgene) virus was delivered to TRAP2 mice (Cat# 030323) of 10-14 weeks of age through stereotactic injection into the LS region (AP: +0.58 mm, ML:0.25 mm, DV:-3.20 mm). Mice were given at least 3 weeks for recovery after surgery. Habituation for i.p. injection was carried out daily for 1 week using injections of 100 µl saline. Mice were i.p. injected with 4-OHT (4-hydroxitamoxifen) at 40 mg/kg and immediately exposed to PO or water in a filter paper in an air-tight mouse cage with familiar bedding in the dark for 5hrs. After cell tagging, animals were kept in a dark and quiet room for 4 additional hours and then returned to their housing room. After 2 weeks, animals were perfused and brains stained for mCherry.

### Stereotaxic injections and Optic Fiber Implantation

Mice were anesthetized with 2% isoflurane, placed in a stereotaxic frame (Kopft Instruments). Eye ointment was applied to the eyes and a subcutaneous injection of meloxicam (2 mg/kg) was given to each mouse after surgery for up to 2 days. Hair was shaved, the scalp was disinfected with iodine solution and an incision was made. The craniotomy was performed using a dental drill (Dremel). A hamilton syringe (Cat# 84851; 85RN; 26s/2”/2) was used to infuse the virus at a rate of 0.1 nL per min at a total volume of 0.5-0.8 uL. After viral infusion, the needle was kept at the injection site for 10 min and then slowly withdrawn. Viral preparations were injected bilaterally or unilaterally at the following coordinates for lateral septum: from bregma, AP: +0.58 mm, ML:0.25 mm, DV:-3.20 mm. For all surgeries, mice were monitored for 72h to ensure full recovery and 2-4 weeks later mice were used in experiments. Optic fiber implants (Doric) for fiber photometry experiments were implanted to the same coordinates as the viral injections with a 0.2 mm difference. Implants were inserted slowly and secured to the mouse skull using two layers of Metabond (Parkell Inc) followed by a layer of dental cement. Mice were single housed and monitored in the first weeks following optic implantation.

### Functional Annotation of RNA Sequencing Data

RNA-seq reads were aligned to the mouse reference genome (mm10) using STAR (v2.7.1a) (60). For each sample, a BAM file was generated containing both mapped and unmapped reads across splice junctions. Secondary alignments and multi-mapped reads were removed using in-house scripts, and only uniquely mapped reads were retained for downstream analyses. GENCODE annotation for mm10 (version 34) was used for reference-guided alignment and downstream quantification. Gene-level expression was quantified using featureCounts (v2.0.1) (61), based on protein-coding genes from the annotation files. Quality control metrics were assessed MultiQC (v1.0.dev0) (62). For differential expression, counts were normalized using counts per million reads (CPM). Genes without reads in either sample were removed. Differential expression analysis was performed in R using a linear model followed by a post-hoc multiple comparison using the R package emmeans. Model comprise P-values were adjusted for multiple comparisons using the Benjamini-Hochberg correction (FDR<0.05). Genes were considered differentially expressed by FDR < 0.05. For GO and enrichment analyses, functional annotation of differentially expressed was performed using scToppR (v0.99.1) (63). Benjamini-Hochberg FDR (FDR<0.05) was applied as a multiple comparison adjustment. Functional categories were filtered based on FDR < 0.05 threshold.

## QUANTIFICATION AND STATISTICAL ANALYSIS

Sample sizes (n = number of animal per group), statistical tests used and statistical significance are described in the Figures and the Figure Legends. Unless otherwise indicated, values are reported as mean ± SEM. (error bars). One-way ANOVA statistical test was followed by post hoc comparisons using Bonferroni multiple comparisons test. Mice with no viral expression were excluded from the final analysis. In figures, asterisks show the obtained statistical significance *p < 0.05, **p < 0.01, ***p < 0.001 and ****p < 0.0001. All statistical analysis was performed using GraphPad Prism 5.0 software (GraphPad).

## DATA AND SOFTWARE AVAILABILITY

The NCBI Gene Expression Omnibus (GEO) accession number for the RNA-seq data (PhosphoTRAP) will be available at GEO. Custom R codes and data to support the analysis, visualizations and functional enrichment analysis are available through GitHub at https://github.com/BioinformaticsMUSC.

## SUPPLEMENTAL INFORMATION

Supplemental information includes 3 figures and 1 table.

## AUTHOR CONTRIBUTIONS

E.P.A. designed the study. E.P.A., M.M., W.S., S.B., S.S. performed and analyzed experiments. P.F., B.D., L.H., A.K., K.., J.C., helped with experiments. J.L., S.B and M.M helped with data analysis and discussion. E.P.A and M.M. wrote the manuscript with input from all authors.

## ACKNOWLEDGMENTS

We thank Rachel Penrod-Martin and the staff of the Animal BioBehavior Core, the veterinary staff at DLAR, the staff of the Genomics Resource Center for technical assistance. This work was funded by the Whitehall Foundation and Medical University of South Carolina funding. SB and SS are supported by the CNDD Genomics and Bioinformatics Core at MUSC (NIH grant P20 GM148302) and by the Biorepository & Tissue Analysis Shared Resource, Hollings Cancer Center, Medical University of South Carolina (P30 CA138313).

## FIGURE LEGENDS

**Supplementary Figure 1. Predator odor exposure alters the translational profile of LS neurons identified by PhosphoTRAP.** Bar plot showing the average log2(IP/INP) values for selected transcripts enriched in PhosphoTRAP immunoprecipitated (IP) samples relative to input (INP) from control and predator odor (PO)-exposed mice. Positive values indicate preferential enrichment of transcripts in activated neurons, whereas negative values indicate relative depletion. Genes displayed (Vdr, Gpr35, Cndp1, Amd1, Ggt1, Ffar3, and Slc38a11) were identified as differentially regulated between PO and control conditions. White bars represent control samples and orange bars represent PO samples. Data are plotted as mean log2(IP/INP), illustrating condition-dependent shifts in translational engagement following acute predator odor stress. All genes showed are significantly different between samples (q value<0.05).

**Supplementary Figure 2. Spatial distribution of marker gene expression in lateral septal clusters from ABC Atlas MERFISH data.** Representative coronal sections from the Allen Brain Cell (ABC) Atlas MERFISH dataset showing imputed spatial expression patterns of selected genes across the lateral septum (LS). The schematic indicates the anatomical location of the LS within the coronal plane. Gene expression maps are displayed for Slc32a1 (GABAergic marker), Crhr2, Glp1r, Oprm1, Nts, Met, and Pax6. Color intensity reflects relative transcript abundance (yellow = lower expression; purple = higher expression).

**Supplementary Figure 3. Ntsr1 expression in hypothalamic regions relevant to LS circuitry.** Coronal schematics (left) indicate the anatomical location of regions analyzed for Ntsr1 expression (black boxes). Representative in situ hybridization images (right) show Ntsr1 mRNA distribution in the lateral hypothalamus (top) and supramammillary nucleus (bottom). Ntsr1-positive cells are detected in both regions, with discrete clusters of labeled neurons in the supramammillary nucleus and a more dispersed pattern in the lateral hypothalamus. These data provide anatomical evidence for Ntsr1-expressing neuronal populations in hypothalamic areas positioned to interact with septal circuits, supporting their potential involvement in neurotensin-related modulation of stress and motivational behaviors.

## REFERENCES

1. Faure, P. (2025). Exploration and behavioral variability. In Handbook of Behavioral Neuroscience, Volume 32, S.J. Cragg and M.E. Walton, eds. (Elsevier), pp. 357–365. 10.1016/B978-0-443-29867-7.00026-8.

2. Thompson SM, Berkowitz LE, Clark BJ. Behavioral and Neural Subsystems of Rodent Exploration. Learn Motiv. 2018 Feb;61:3–15. doi: 10.1016/j.lmot.2017.03.009. Epub 2017 Apr 13. PMID: 30270939; PMCID: PMC6159932.

3. Perusini JN, Fanselow MS. Neurobehavioral perspectives on the distinction between fear and anxiety. Learn Mem. 2015 Aug 18;22(9):417–25. doi: 10.1101/lm.039180.115. PMID: 26286652; PMCID: PMC4561408.

4. McManus, H., Milad, M.R. (2025). Behavioral and Brain Mechanisms of Active Avoidance and Their Relevance to Anxiety Disorders. In: Blackford, J.U., Milad, M.R. (eds) New Discoveries in the Brain Sciences of Fear and Anxiety - From Basic to Clinical Neuroscience. Current Topics in Behavioral Neurosciences, vol 73. Springer, Cham. 10.1007/7854_2025_593

5. López-Moraga A, Beckers T, Luyten L. The effects of stress on avoidance in rodents: An unresolved matter. Front Behav Neurosci. 2022 Sep 28;16:983026. doi: 10.3389/fnbeh.2022.983026. PMID: 36275848; PMCID: PMC9580497.

6. McEwen BS, Akil H. Revisiting the Stress Concept: Implications for Affective Disorders. J Neurosci. 2020 Jan 2;40(1):12–21. doi: 10.1523/JNEUROSCI.0733-19.2019. PMID: 31896560; PMCID: PMC6939488.

7. McEwen BS, Bowles NP, Gray JD, Hill MN, Hunter RG, Karatsoreos IN, Nasca C. Mechanisms of stress in the brain. Nat Neurosci. 2015 Oct;18(10):1353–63. doi: 10.1038/nn.4086. Epub 2015 Sep 25. PMID: 26404710; PMCID: PMC4933289.

8. Verbitsky A, Dopfel D, Zhang N. Rodent models of post-traumatic stress disorder: behavioral assessment. Transl Psychiatry. 2020 May 6;10(1):132. doi: 10.1038/s41398-020-0806-x. PMID: 32376819; PMCID: PMC7203017.

9. Kigar SL, Cuarenta A, Zuniga CL, Chang L, Auger AP, Bakshi VP. Brain, behavior, and physiological changes associated with predator stress-An animal model for trauma exposure in adult and neonatal rats. Front Mol Neurosci. 2024 Feb 29;17:1322273. doi: 10.3389/fnmol.2024.1322273. PMID: 38486962; PMCID: PMC10938396.

10. Adamec R. Transmitter systems involved in neural plasticity underlying increased anxiety and defense--implications for understanding anxiety following traumatic stress. Neurosci Biobehav Rev. 1997 Nov;21(6):755–65. doi: 10.1016/s0149-7634(96)00055-3. PMID: 9415900.

11. Adamec RE, Blundell J, Burton P. Phosphorylated cyclic AMP response element binding protein expression induced in the periaqueductal gray by predator stress: its relationship to the stress experience, behavior and limbic neural plasticity. Prog Neuropsychopharmacol Biol Psychiatry. 2003 Dec;27(8):1243–67. doi: 10.1016/j.pnpbp.2003.09.017. PMID: 14659479.

12. Baumbach JL, Mui CYY, Leonetti AM, Martin LJ. Corticosterone regulates the balance between freezing and rearing in defensive responses to predator threat. Prog Neuropsychopharmacol Biol Psychiatry. 2026 Jan 2;144:111579. doi: 10.1016/j.pnpbp.2025.111579. Epub 2025 Dec 8. PMID: 41371387.

13. Pichlmeier O, Caner E, von Kalben L, Grodzki LM, Weigelt N, Pepe C, Morellini F. Reconsolidation of episodic-like memory is affected by exposure to the predator odor TMT in mice. Behav Brain Res. 2025 Oct 18;495:115817. doi: 10.1016/j.bbr.2025.115817. Epub 2025 Sep 10. PMID: 40939869.

14. Xu H, Yuan H, Song H, Pan Y, Wu Z, Wang S, Cheng J, Luo Y, Lin L, Min P, Yue Y, Chen X, Zhang K, Fukunaga K, Sasaki T, Mao X, Han F, Lu YM. Multimode neural population coding of diverse innate fear response by mitral and tufted cells. Cell Rep. 2025 Sep 23;44(9):116255. doi: 10.1016/j.celrep.2025.116255. Epub 2025 Sep 8. PMID: 40924580.

15. Ivanova D, Voliotis M, Tsaneva-Atanasova K, O’Byrne KT, Li XF. NK3R signalling in the posterodorsal medial amygdala is involved in stress-induced suppression of pulsatile LH secretion in female mice. J Neuroendocrinol. 2024 May;36(5):e13384. doi: 10.1111/jne.13384. Epub 2024 Mar 22. PMID: 38516965; PMCID: PMC11411622.

16. Genné-Bacon EA, Trinko JR, DiLeone RJ. Innate Fear-Induced Weight Regulation in the C57BL/6J Mouse. Front Behav Neurosci. 2016 Jul 4;10:132. doi: 10.3389/fnbeh.2016.00132. PMID: 27458352; PMCID: PMC4930939.

17. Kaegi ZE, Carter ME. Tachykinin-1-expressing parasubthalamic nucleus neurons are necessary for odorant-induced appetite suppression. Physiol Behav. 2025 Apr 1;292:114836. doi: 10.1016/j.physbeh.2025.114836. Epub 2025 Jan 31. PMID: 39892639; PMCID: PMC11846689.\

18. Kondev V, Morgan A, Najeed M, Winters ND, Kingsley PJ, Marnett L, Patel S. The Endocannabinoid 2-Arachidonoylglycerol Bidirectionally Modulates Acute and Protracted Effects of Predator Odor Exposure. Biol Psychiatry. 2022 Nov 1;92(9):739–749. doi: 10.1016/j.biopsych.2022.05.012. Epub 2022 May 17. PMID: 35961791; PMCID: PMC9827751.

19. Saito H, Nishizumi H, Suzuki S, Matsumoto H, Ieki N, Abe T, Kiyonari H, Morita M, Yokota H, Hirayama N, Yamazaki T, Kikusui T, Mori K, Sakano H. Immobility responses are induced by photoactivation of single glomerular species responsive to fox odour TMT. Nat Commun. 2017 Jul 7;8:16011. doi: 10.1038/ncomms16011. PMID: 28685774; PMCID: PMC5504302.

20. Takahashi LK. Olfactory systems and neural circuits that modulate predator odor fear. Front Behav Neurosci. 2014 Mar 11;8:72. doi: 10.3389/fnbeh.2014.00072. PMID: 24653685; PMCID: PMC3949219.

21. Zak JD, Reddy G, Konanur V, Murthy VN. Distinct information conveyed to the olfactory bulb by feedforward input from the nose and feedback from the cortex. Nat Commun. 2024 Apr 16;15(1):3268. doi: 10.1038/s41467-024-47366-6. PMID: 38627390; PMCID: PMC11021479.

22. Smith W, Azevedo EP. Hunger Games: A Modern Battle Between Stress and Appetite. J Neurochem. 2025 Feb;169(2):e70006. doi: 10.1111/jnc.70006. PMID: 39936619.

23. Azevedo EP, Tan B, Pomeranz LE, Ivan V, Fetcho R, Schneeberger M, Doerig KR, Liston C, Friedman JM, Stern SA. A limbic circuit selectively links active escape to food suppression. Elife. 2020 Sep 7;9:e58894. doi: 10.7554/eLife.58894. PMID: 32894221; PMCID: PMC7476759.

24. Bhatti Mazo DL, Berger MZC, Pasqualini AL, Wu SJ, Reid CM, Brito SI, Qiu S, Levitt P, Harwell CC, Anthony TE, Fishell G. Feature-specific threat coding in lateral septum guides defensive action. Nature. 2026 May 20. doi: 10.1038/s41586-026-10520-9. Epub ahead of print. Erratum in: Nature. 2026 Jul 24. doi: 10.1038/s41586-026-10911-y. PMID: 42162422.

25. García MT, Tran DN, Peterson RE, Stegmann SK, Hanson SM, Reid CM, Xie Y, Vu S, Harwell CC. A developmentally defined population of neurons in the lateral septum controls responses to aversive stimuli. bioRxiv [Preprint]. 2023 Oct 9:2023.09.24.559205. doi: 10.1101/2023.09.24.559205. PMID: 37873286; PMCID: PMC10592641.

26. Chen M, Li J, Shan W, Yang J, Zuo Z. Auditory fear memory retrieval requires BLA-LS and LS-VMH circuitries via GABAergic and dopaminergic neurons. EMBO Rep. 2025 Apr;26(7):1816–1834. doi: 10.1038/s44319-025-00403-x. Epub 2025 Mar 7. PMID: 40055468; PMCID: PMC11977213.

27. Patel H. The role of the lateral septum in neuropsychiatric disease. J Neurosci Res. 2022 Jul;100(7):1422–1437. doi: 10.1002/jnr.25052. Epub 2022 Apr 20. PMID: 35443088.

28. Randler C, Kalb J. Predator avoidance behavior of nocturnal and diurnal rodents. Behav Processes. 2020 Oct;179:104214. doi: 10.1016/j.beproc.2020.104214. Epub 2020 Aug 6. PMID: 32768461.

29. McEwen BS. Allostasis and the Epigenetics of Brain and Body Health Over the Life Course: The Brain on Stress. JAMA Psychiatry. 2017 Jun 1;74(6):551–552. doi: 10.1001/jamapsychiatry.2017.0270. PMID: 28445556.

30. Knight ZA, Tan K, Birsoy K, Schmidt S, Garrison JL, Wysocki RW, Emiliano A, Ekstrand MI, Friedman JM. Molecular profiling of activated neurons by phosphorylated ribosome capture. Cell. 2012 Nov 21;151(5):1126–37. doi: 10.1016/j.cell.2012.10.039. PMID: 23178128; PMCID: PMC3839252.

31. Simon RC, Fleming WT, Briones BA, Trzeciak M, Senthilkumar P, Ishii KK, Hjort MM, Martin MM, Hashikawa K, Sanders AD, Golden SA, Stuber GD. Opioid-driven disruption of the septum reveals a role for neurotensin-expressing neurons in withdrawal. Neuron. 2025 Jul 23;113(14):2325–2343.e9. doi: 10.1016/j.neuron.2025.04.024. Epub 2025 May 15. PMID: 40378834.

32. Liebelt F, Vertegaal AC. Ubiquitin-dependent and independent roles of SUMO in proteostasis. Am J Physiol Cell Physiol. 2016 Aug 1;311(2):C284–96. doi: 10.1152/ajpcell.00091.2016. Epub 2016 Jun 22. PMID: 27335169; PMCID: PMC5129774.

33. He J, Zhang P, Shen L, Niu L, Tan Y, Chen L, Zhao Y, Bai L, Hao X, Li X, Zhang S, Zhu L. Short-Chain Fatty Acids and Their Association with Signalling Pathways in Inflammation, Glucose and Lipid Metabolism. Int J Mol Sci. 2020 Sep 2;21(17):6356. doi: 10.3390/ijms21176356. PMID: 32887215; PMCID: PMC7503625.

34. Mitrić A, Castellano I. Targeting gamma-glutamyl transpeptidase: A pleiotropic enzyme involved in glutathione metabolism and in the control of redox homeostasis. Free Radic Biol Med. 2023 Nov 1;208:672–683. doi: 10.1016/j.freeradbiomed.2023.09.020. Epub 2023 Sep 20. PMID: 37739139.

35. Zhou L, Zhang S, Zhang Y, Luo Y, Sun X. Carnosine Dipeptidase(Cndp): An emerging therapeutic target for metabolic diseases and cancers. Genes Dis. 2025 Aug 13;13(1):101804. doi: 10.1016/j.gendis.2025.101804. PMID: 41098976; PMCID: PMC12519243.

36. Zabala-Letona A, Arruabarrena-Aristorena A, Martín-Martín N, Fernandez-Ruiz S, Sutherland JD, Clasquin M, Tomas-Cortazar J, Jimenez J, Torres I, Quang P, Ximenez-Embun P, Bago R, Ugalde-Olano A, Loizaga-Iriarte A, Lacasa-Viscasillas I, Unda M, Torrano V, Cabrera D, van Liempd SM, Cendon Y, Castro E, Murray S, Revandkar A, Alimonti A, Zhang Y, Barnett A, Lein G, Pirman D, Cortazar AR, Arreal L, Prudkin L, Astobiza I, Valcarcel-Jimenez L, Zuñiga-García P, Fernandez-Dominguez I, Piva M, Caro-Maldonado A, Sánchez-Mosquera P, Castillo-Martín M, Serra V, Beraza N, Gentilella A, Thomas G, Azkargorta M, Elortza F, Farràs R, Olmos D, Efeyan A, Anguita J, Muñoz J, Falcón-Pérez JM, Barrio R, Macarulla T, Mato JM, Martinez-Chantar ML, Cordon-Cardo C, Aransay AM, Marks K, Baselga J, Tabernero J, Nuciforo P, Manning BD, Marjon K, Carracedo A. mTORC1-dependent AMD1 regulation sustains polyamine metabolism in prostate cancer. Nature. 2017 Jul 6;547(7661):109–113. doi: 10.1038/nature22964. Epub 2017 Jun 28. Erratum in: Nature. 2018 Feb 22;554(7693):554. doi: 10.1038/nature25470. PMID: 28658205; PMCID: PMC5505479.

37. Angelucci F, Cerman J, Amlerova J, Sheardova K, Pavlik J, Hort J. Spermidine in Alzheimer’s Disease: Evidence from Animal Models and Human Studies. Degener Neurol Neuromuscul Dis. 2026 Jun 20;16:608341. doi: 10.2147/DNND.S608341. PMID: 42358231; PMCID: PMC13292852.

38. Bröer S. The SLC38 family of sodium-amino acid co-transporters. Pflugers Arch. 2014 Jan;466(1):155–72. doi: 10.1007/s00424-013-1393-y. Epub 2013 Nov 6. PMID: 24193407.

39. Voltan G, Cannito M, Ferrarese M, Ceccato F, Camozzi V. Vitamin D: An Overview of Gene Regulation, Ranging from Metabolism to Genomic Effects. Genes (Basel). 2023 Aug 25;14(9):1691. doi: 10.3390/genes14091691. PMID: 37761831; PMCID: PMC10531002.

40. Wang J, Simonavicius N, Wu X, Swaminath G, Reagan J, Tian H, Ling L. Kynurenic acid as a ligand for orphan G protein-coupled receptor GPR35. J Biol Chem. 2006 Aug 4;281(31):22021–22028. doi: 10.1074/jbc.M603503200. Epub 2006 Jun 5. PMID: 16754668.

41. Zhang M, Pan X, Jung W, Halpern AR, Eichhorn SW, Lei Z, Cohen L, Smith KA, Tasic B, Yao Z, Zeng H, Zhuang X. Molecularly defined and spatially resolved cell atlas of the whole mouse brain. Nature. 2023 Dec;624(7991):343–354. doi: 10.1038/s41586-023-06808-9. Epub 2023 Dec 13. PMID: 38092912; PMCID: PMC10719103.

42. Yao Z, van Velthoven CTJ, Kunst M, Zhang M, McMillen D, Lee C, Jung W, Goldy J, Abdelhak A, Aitken M, Baker K, Baker P, Barkan E, Bertagnolli D, Bhandiwad A, Bielstein C, Bishwakarma P, Campos J, Carey D, Casper T, Chakka AB, Chakrabarty R, Chavan S, Chen M, Clark M, Close J, Crichton K, Daniel S, DiValentin P, Dolbeare T, Ellingwood L, Fiabane E, Fliss T, Gee J, Gerstenberger J, Glandon A, Gloe J, Gould J, Gray J, Guilford N, Guzman J, Hirschstein D, Ho W, Hooper M, Huang M, Hupp M, Jin K, Kroll M, Lathia K, Leon A, Li S, Long B, Madigan Z, Malloy J, Malone J, Maltzer Z, Martin N, McCue R, McGinty R, Mei N, Melchor J, Meyerdierks E, Mollenkopf T, Moonsman S, Nguyen TN, Otto S, Pham T, Rimorin C, Ruiz A, Sanchez R, Sawyer L, Shapovalova N, Shepard N, Slaughterbeck C, Sulc J, Tieu M, Torkelson A, Tung H, Valera Cuevas N, Vance S, Wadhwani K, Ward K, Levi B, Farrell C, Young R, Staats B, Wang MM, Thompson CL, Mufti S, Pagan CM, Kruse L, Dee N, Sunkin SM, Esposito L, Hawrylycz MJ, Waters J, Ng L, Smith K, Tasic B, Zhuang X, Zeng H. A high-resolution transcriptomic and spatial atlas of cell types in the whole mouse brain. Nature. 2023 Dec;624(7991):317–332. doi: 10.1038/s41586-023-06812-z. Epub 2023 Dec 13. PMID: 38092916; PMCID: PMC10719114.

43. Park S, Guo Y, Negre J, Preto J, Smithers CC, Azad AK, Overduin M, Murray AG, Eitzen G. Fgd5 is a Rac1-specific Rho GEF that is selectively inhibited by aurintricarboxylic acid. Small GTPases. 2021 Mar;12(2):147–160. doi: 10.1080/21541248.2019.1674765. Epub 2019 Oct 10. PMID: 31601145; PMCID: PMC7849785.

44. Shang C, Chen Z, Liu A, Li Y, Zhang J, Qu B, Yan F, Zhang Y, Liu W, Liu Z, Guo X, Li D, Wang Y, Cao P. Divergent midbrain circuits orchestrate escape and freezing responses to looming stimuli in mice. Nat Commun. 2018 Mar 26;9(1):1232. doi: 10.1038/s41467-018-03580-7. PMID: 29581428; PMCID: PMC5964329.

45. DeNardo LA, Liu CD, Allen WE, Adams EL, Friedmann D, Fu L, Guenthner CJ, Tessier-Lavigne M, Luo L. Temporal evolution of cortical ensembles promoting remote memory retrieval. Nat Neurosci. 2019 Mar;22(3):460–469. doi: 10.1038/s41593-018-0318-7. Epub 2019 Jan 28. PMID: 30692687; PMCID: PMC6387639.

46. Goode TD, Bernstein MX, Totty MS, Alipio JB, Vicidomini C, Pathak D, Besnard A, Chizari D, Sachdev N, Kritzer MD, Chung A, Duan X, Macosko E, Hicks SC, Zweifel LS, Sahay A. A dorsal hippocampus-prodynorphinergic dorsolateral septum-to-lateral hypothalamus circuit mediates contextual gating of feeding. Neuron. 2026 Jun 3;114(11):2050–2072.e14. doi: 10.1016/j.neuron.2026.01.025. Epub 2026 Feb 12. PMID: 41687615; PMCID: PMC13288035.

47. Allen Institute (2004). Allen Mouse Brain Atlas [dataset 73519704]. Available from mouse.brain-map.org. Allen Institute for Brain Science (2011).

48. Fuchs E, Flugge G, Czeh B. Remodeling of neuronal networks by stress. Front Biosci. 2006 Sep 1;11:2746–58. doi: 10.2741/2004. PMID: 16720347.

49. Rossi MJ, Pekkurnaz G. Powerhouse of the mind: mitochondrial plasticity at the synapse. Curr Opin Neurobiol. 2019 Aug;57:149–155. doi: 10.1016/j.conb.2019.02.001. Epub 2019 Mar 12. PMID: 30875521; PMCID: PMC6629504.

50. Tanaka K, Masu M, Nakanishi S. Structure and functional expression of the cloned rat neurotensin receptor. Neuron. 1990 Jun;4(6):847–54. doi: 10.1016/0896-6273(90)90137-5. PMID: 1694443.

51. Leinninger GM, Opland DM, Jo YH, Faouzi M, Christensen L, Cappellucci LA, Rhodes CJ, Gnegy ME, Becker JB, Pothos EN, Seasholtz AF, Thompson RC, Myers MG Jr. Leptin action via neurotensin neurons controls orexin, the mesolimbic dopamine system and energy balance. Cell Metab. 2011 Sep 7;14(3):313–23. doi: 10.1016/j.cmet.2011.06.016. PMID: 21907138; PMCID: PMC3183584.

52. Muschol M, Salzberg BM. Dependence of transient and residual calcium dynamics on action-potential patterning during neuropeptide secretion. J Neurosci. 2000 Sep 15;20(18):6773–80. doi: 10.1523/JNEUROSCI.20-18-06773.2000. PMID: 10995820; PMCID: PMC6772822.

53. Kitabgi P, De Nadai F, Cuber JC, Dubuc I, Nouel D, Costentin J. Calcium-dependent release of neuromedin N and neurotensin from mouse hypothalamus. Neuropeptides. 1990 Feb;15(2):111–4. doi: 10.1016/0143-4179(90)90047-3. PMID: 2080018.

54. Tabarean IV. Opposing actions of co-released GABA and neurotensin on the activity of preoptic neurons and on body temperature. Elife. 2024 Aug 29;13:RP98677. doi: 10.7554/eLife.98677. PMID: 39207910; PMCID: PMC11361704.

55. NIMH. (2017). Any Anxiety Disorder.

56. Hao Y, Hao S, Andersen-Nissen E, Mauck WM 3rd, Zheng S, Butler A, Lee MJ, Wilk AJ, Darby C, Zager M, Hoffman P, Stoeckius M, Papalexi E, Mimitou EP, Jain J, Srivastava A, Stuart T, Fleming LM, Yeung B, Rogers AJ, McElrath JM, Blish CA, Gottardo R, Smibert P, Satija R. Integrated analysis of multimodal single-cell data. Cell. 2021 Jun 24;184(13):3573–3587.e29. doi: 10.1016/j.cell.2021.04.048. Epub 2021 May 31. PMID:34062119; PMCID: PMC8238499.

57. Germain PL, Lun A, Garcia Meixide C, Macnair W, Robinson MD. Doublet identification in single-cell sequencing data using *scDblFinder*. F1000Res. 2021 Sep 28;10:979. doi: 10.12688/f1000research.73600.2. PMID: 35814628; PMCID: PMC9204188.

58. Hafemeister C, Satija R. Normalization and variance stabilization of single-cell RNA-seq data using regularized negative binomial regression. Genome Biol. 2019 Dec 23;20(1):296. doi: 10.1186/s13059-019-1874-1. PMID: 31870423; PMCID: PMC6927181.

59. Fan J, Lyu Y, Zhang Q, Wang X, Li M, Xiao R. MuSiC2: cell-type deconvolution for multi-condition bulk RNA-seq data. Brief Bioinform. 2022 Nov 19;23(6):bbac430. doi: 10.1093/bib/bbac430. PMID: 36208175; PMCID: PMC9677503.

60. Dobin A, Davis CA, Schlesinger F, Drenkow J, Zaleski C, Jha S, Batut P, Chaisson M, Gingeras TR. STAR: ultrafast universal RNA-seq aligner. Bioinformatics. 2013 Jan 1;29(1):15–21. doi: 10.1093/bioinformatics/bts635. Epub 2012 Oct 25. PMID: 23104886; PMCID: PMC3530905.

61. Liao Y, Smyth GK, Shi W. featureCounts: an efficient general purpose program for assigning sequence reads to genomic features. Bioinformatics. 2014 Apr 1;30(7):923–30. doi: 10.1093/bioinformatics/btt656. Epub 2013 Nov 13. PMID: 24227677.

62. Ewels P, Magnusson M, Lundin S, Käller M. MultiQC: summarize analysis results for multiple tools and samples in a single report. Bioinformatics. 2016 Oct 1;32(19):3047–8. doi: 10.1093/bioinformatics/btw354. Epub 2016 Jun 16. PMID: 27312411; PMCID: PMC5039924.

63. Granger B, Berto S. scToppR: a coding-friendly R interface to ToppGene. Bioinformatics. 2024 Nov 1;40(11):btae582. doi: 10.1093/bioinformatics/btae582. PMID: 39340795; PMCID: PMC11552619.

