## Supplemental Figures for "A Neurotensin Brake on Exploratory Drive under Persistent Threat"

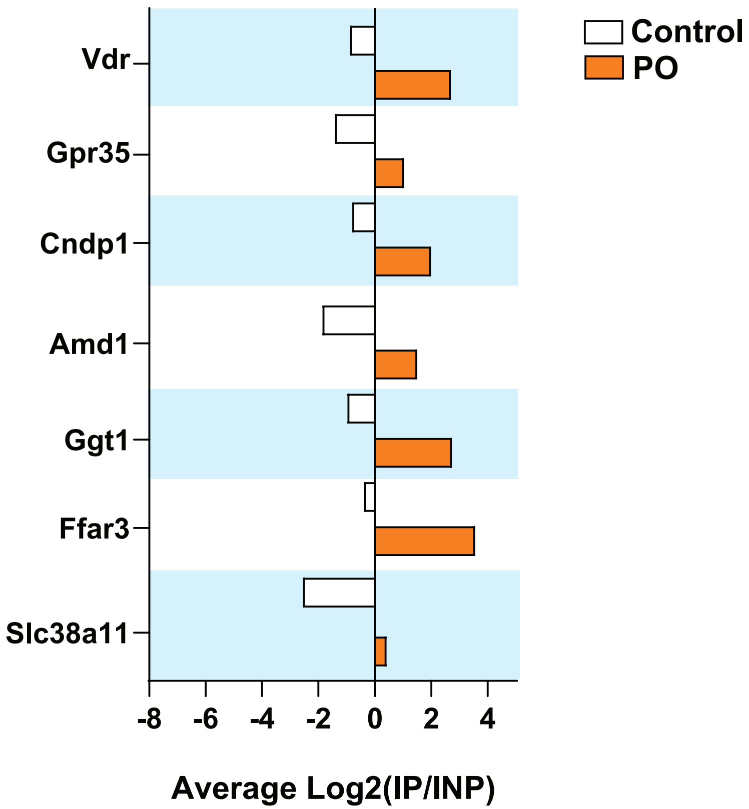


Supplementary Fig.1

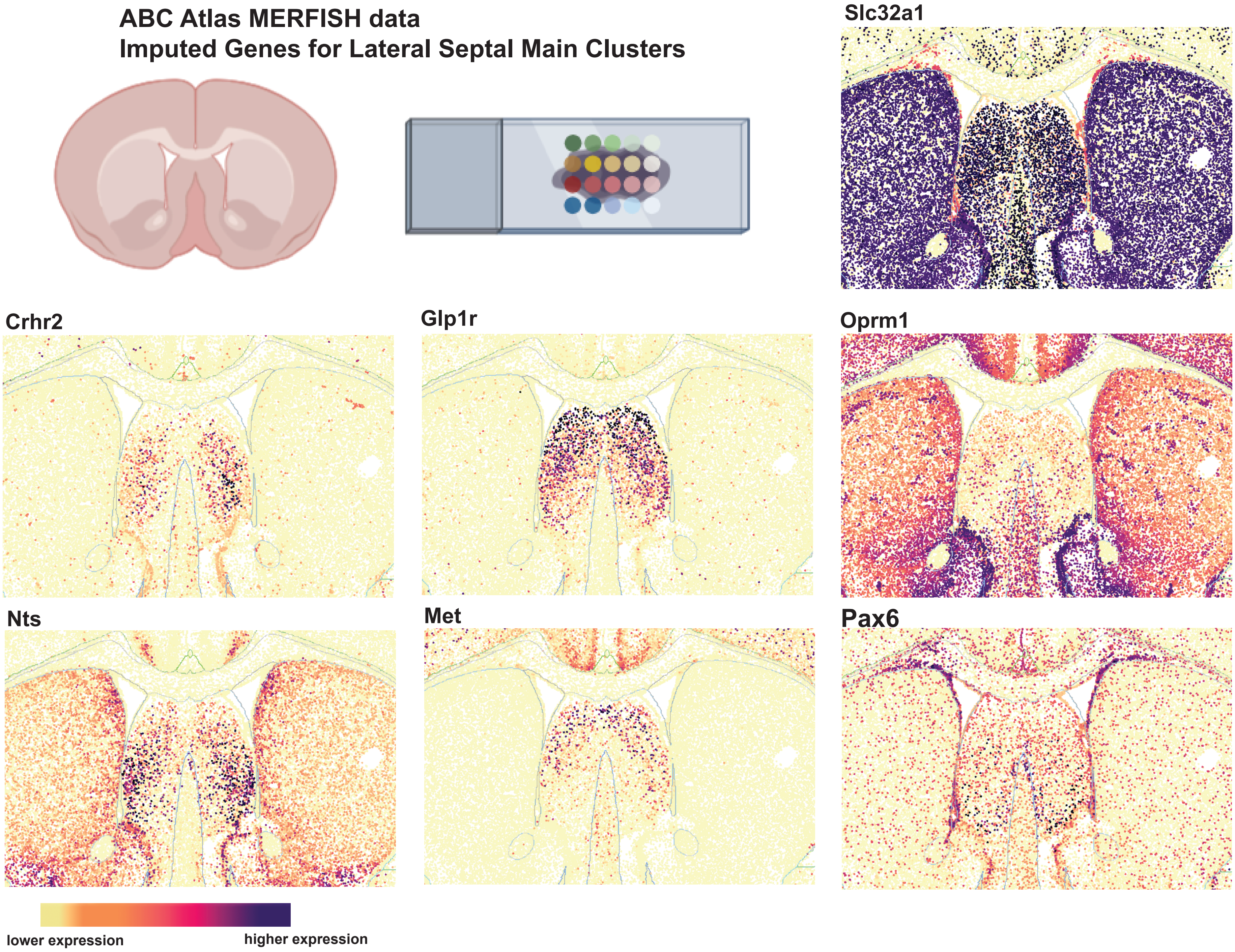


Supplementary Fig.2

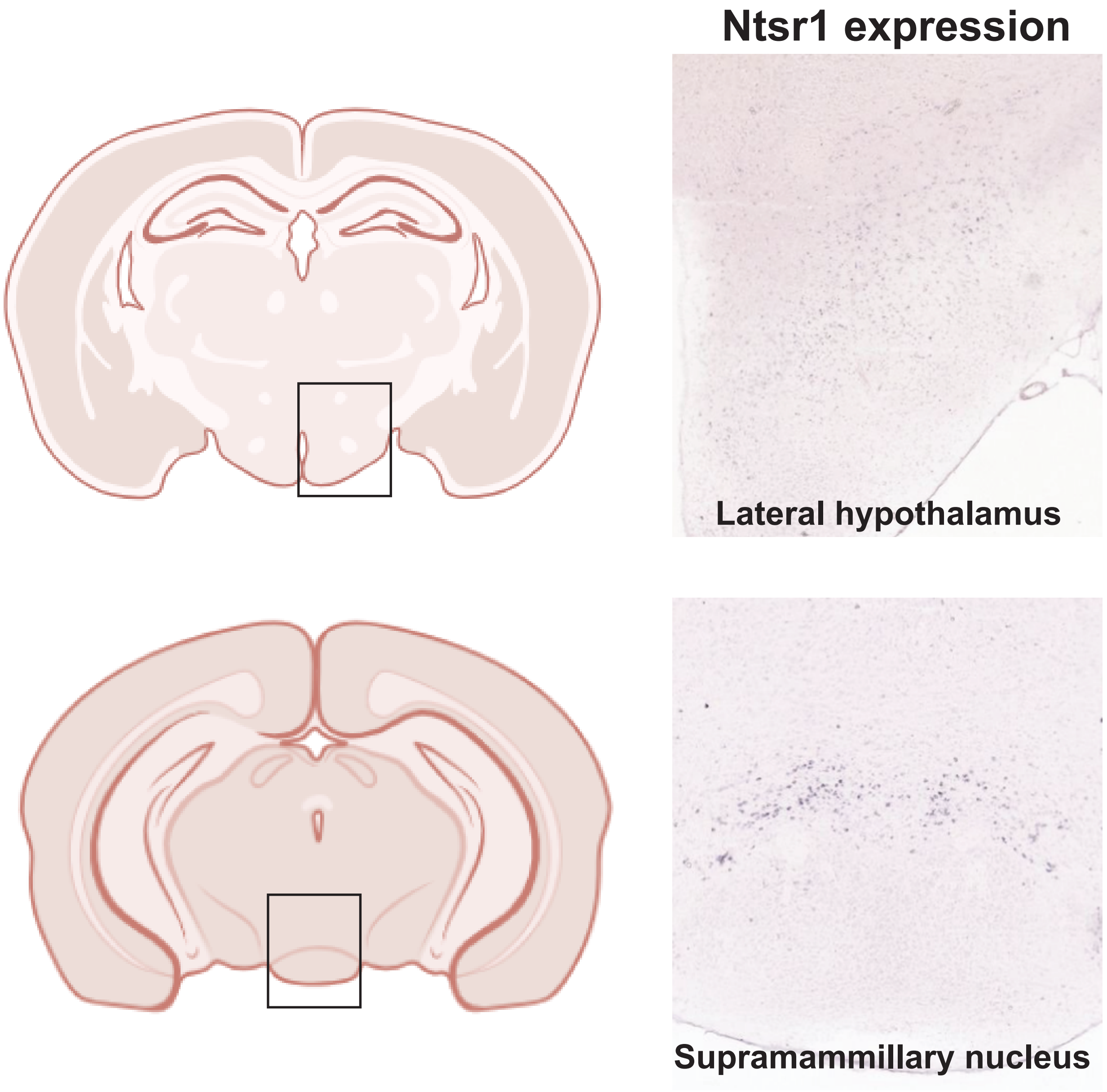


Supplementary Fig.3

Supplementary Table 1. Whole Brain Fos mapping

| Brain Region | Abbreviation | Control (Mean ± SEM) | PO (Mean ± SEM) | p-value | Significance |
|---|---|---|---|---|---|
| Piriform Cortex | Pir | 4.67 ± 2.73 | 76.67 ± 45.67 | 0.190666 | ns |
| Bed nucleus of stria terminalis | BNST | 1.33 ± 0.67 | 38.67 ± 22.26 | 0.168947 | ns |
| Insular Cortex | IC | 8.33 ± 0.88 | 81.67 ± 20.69 | 0.023991 | * |
| Claustrum | Cla | 10.67 ± 4.33 | 51.33 ± 15.45 | 0.064392 | ns |
| Lateral Septum | LS | 4.33 ± 2.85 | 207 ± 70.15 | 0.044712 | * |
| Paraventricular Nucleus of the Hypothalamus | PVH | 2 ± 1 | 116 ± 38.31 | 0.040967 | * |
| 7n | 7n | 7.33 ± 2.6 | 45 ± 14.05 | 0.057793 | ns |
| Superior Colliculus | SC | 97.67 ± 17.98 | 218.67 ± 36.73 | 0.041585 | * |
| Gi | Gi | 29 ± 9.02 | 38.33 ± 11.61 | 0.560001 | ns |
| Lateral Entorhinal Cortex | LEC | 4.33 ± 0.67 | 61.67 ± 28.67 | 0.116188 | ns |
| Medial Preoptic Area | MPO | 61.33 ± 14.84 | 152.67 ± 23.67 | 0.030815 | * |
| Periaqueductal Gray | PAG | 53.67 ± 21.87 | 124.33 ± 6.77 | 0.036666 | * |
| Parabrachial Nucleus | PBN | 12.67 ± 2.19 | 74.33 ± 6.98 | 0.001086 | ** |
| Pontine | Pontine | 16.67 ± 8.09 | 193 ± 46.76 | 0.020547 | * |
| PVA | PVA | 51 ± 21.7 | 316.33 ± 109.55 | 0.076327 | ns |
| Paraventricular Thalamus | PVT | 35.67 ± 13.13 | 139 ± 20.53 | 0.013257 | * |
| Lateral Habenula | LHb | 5.67 ± 4.18 | 36.33 ± 15.34 | 0.126044 | ns |
| RMg | RMg | 9.33 ± 3.28 | 47 ± 15.62 | 0.077679 | ns |
| RPA | RPA | 3.67 ± 1.67 | 17 ± 1.53 | 0.004135 | ** |

| | | | | | |
|---|---|---|---|---|---|
| Hippocampus | Hip | 136 ± 21.73 | 139.67 ± 16.9 | 0.900490 | ns |
| Ventral Tegmental Area | VTA | 35.67 ± 23.25 | 134 ± 18.08 | 0.028887 | * |
| Central Amygdala | CeA | 9.67 ± 1.86 | 30.33 ± 13.09 | 0.193147 | ns |
| Basolateral Amygdala | BLA | 23 ± 2.31 | 99.33 ± 1.86 | 0.017995 | * |
| Cingulate Cortex | Cing | 16 ± 6.43 | 196.33 ± 21.61 | 0.001325 | ** |
| Lateral Hypothalamic Area | LHA | 27.33 ± 3.38 | 91 ± 23.52 | 0.055240 | ns |
| Supramammillary Nucleus | SUM | 26 ± 8.54 | 106.33 ± 25.73 | 0.041407 | * |
